# Drug screening at single-organoid resolution via bioprinting and interferometry

**DOI:** 10.1101/2021.10.03.462896

**Authors:** Peyton J. Tebon, Bowen Wang, Alexander L. Markowitz, Ardalan Davarifar, Patrycja Krawczuk, Graeme Murray, Huyen Thi Lam Nguyen, Nasrin Tavanaie, Thang L. Nguyen, Paul C. Boutros, Michael A. Teitell, Alice Soragni

## Abstract

High-throughput drug screening is an established approach to investigate tumor biology and identify therapeutic leads. Traditional platforms for high-throughput screening use two-dimensional cultures of immortalized cell lines which do not accurately reflect the biology of human tumors. More clinically relevant model systems, such as three-dimensional tumor organoids, can be difficult to screen and scale. For example, manually seeded organoids coupled to destructive endpoint assays allow for the characterization of response to treatment, but do not capture the transitory changes and intra-sample heterogeneity underlying clinically observed resistance to therapy. We therefore developed a pipeline to generate bioprinted tumor organoids linked to label-free, real-time imaging *via* high-speed live cell interferometry (HSLCI) and machine learning-based quantitation of individual organoids. Bioprinting cells gives rise to 3D organoid structures that preserve tumor histology and gene expression. HSLCI imaging in tandem with machine learning-based image segmentation and organoid classification tools enables accurate, label-free parallel mass measurements for thousands of bioprinted organoids. We demonstrate that our method quantitatively identifies individual organoids as insensitive, transiently sensitive, or persistently sensitive to specific treatments. This opens new avenues for rapid, actionable therapeutic selection using automated tumor organoid screening.

## Introduction

Functional precision oncology involves exposing tumor cells from individual patients to candidate therapeutic interventions ex vivo^1–4^. By monitoring response, treatment regimens with a higher probability of success can be prioritized^5–9^. These types of assays can provide useful sensitivity profiles even for tumors that lack currently actionable genomic alterations, and are thus incompatible with genomic-based precision medicine approaches^10^, By directly measuring the effect of drugs on tissues or cells, functional assays can inform on the therapeutic resistance and sensitivity landscape of tumors without requiring full knowledge of the underlying molecular vulnerabilities a priori^11,12^.

The broadly adopted model systems used in screening assays to identify possible leads all have limitations. Two-dimensional cell lines are relatively simple and inexpensive to culture but fail to represent the architecture, behavior and drug response of native tissue^13,14^. Mouse models have additional complexity but carry inherent, species-specific variations that limit their translation to human patients^15^. Patient-derived xenograft (PDX) models aim to better recapitulate human cancers yet are constrained by the large cost and time associated with their use, making large drug screening studies practically challenging^16^. Three-dimensional (3D) tumor organoids are promising models for precision medicine that can be established rapidly and effectively from a variety of cell types and tissue sources, and accurately mimic a patient’s response to therapy^4,6–9,12,17^. They are physiologically-relevant, personalized cancer models well-suited for drug development and clinical applications^18,19^. The key outstanding limitations to the broad adoption of organoid-based screenings remain the time-intensive and operator-to-operator susceptibility of the cell seeding steps as well as destructive, population-level approaches required for subsequent organoid analysis^20^.

To overcome these limitations, we developed an organoid screening pipeline that combines automated cell seeding via bioprinting with high-speed live cell interferometry (HSLCI) and machine learning-based image segmentation and classification and for non-invasive, label-free, real-time organoid imaging. The new pipeline is based on our previously developed screening approach that takes advantage of patient-derived tumor organoids seeded in a mini-ring format to automate high-throughput drug testing, with results available within one week from surgery^6,7,21^. We automate cell seeding by including bioprinting, a technique for precise, reproducible deposition of cells in bioinks onto solid supports, to seed the organoids^22^. Bioprinting has rapidly gained traction in cancer biology as embedded cells can interact with physiological microenvironment components in the bioink to create physiologically-representative tumor models^20,22–26^.

We then implement HSLCI to rapidly monitor changes in dry biomass and biomass distribution of single organoids over time. HSLCI, a type of quantitative phase imaging (QPI)^27–32^, measures the phase shift of light transmitted through the sample using a wavefront sensing camera^30,33^. Due to the defined linear relationship between the refractive index and mass density and refractive index of biomolecules in solution, which is invariant with respect to changes in cellular content^34–38^, measured phase shifts can be integrated across the area of an image and multiplied by a conversion factor to obtain the dry biomass density of imaged cells^33^. Biomass is an important metric of organoid fitness as its dynamics are the direct result of biosynthetic and degradative processes within cells^33^. In previous work, QPI measurements of biomass changes allowed resolution of drug-resistant and drug-sensitive cells in 2D cell culture models within hours of treatment^27–29,31,32,39,40^. HSLCI-measured response profiles have also been shown to match drug sensitivity from patient-derived xenograft (PDX) mouse models of breast cancer^28^. However, HSLCI has been applied exclusively to screening cancer lines grown in 2D or single-cell suspensions of excised PDX tumors thus far^27–32^. We demonstrate here that bioprinted organoids deposited in uniform, flat layers of extracellular matrix allow label-free, real-time, non-destructive quantification of growth patterns and drug responses at single-organoid resolution.

## Results

### Bioprinting enables seeding cells in Matrigel in suitable geometries for quantitative imaging applications

To address current limitations^6,7,21^ and facilitate non-invasive, label-free, real time imaging of 3D organoids by HSLCI, we created an automated cell printing protocol using an extrusion bioprinter. As a base, we used an organoid platform that seeds cells in mini-rings of Matrigel around the rim of 96-well plates, with the empty center allowing the use of automated liquid handlers, facilitating media exchanges and addition of perturbagens^6,7,21^. We retained the empty center architecture but altered the geometry to bioprint mini-squares of cells in Matrigel (**Figure 1A**). Positioning the sides of the square in the HSLCI imaging path allows sampling of a larger area and limits imaging artifacts caused by uneven illumination at well edges^30^ (**Figure 1A**). Our bioprinting protocol entails suspending cells in a bioink consisting of a 3:4 ratio of medium to Matrigel. This material is then transferred to a print cartridge, incubated at 17°C for 30 minutes, and bioprinted into each well at a pressure between 12 and 15 kPa, resulting in ~200 *μ*m prints on standard glass-bottom plates (**Figure 1B**).

**Figure 1.**
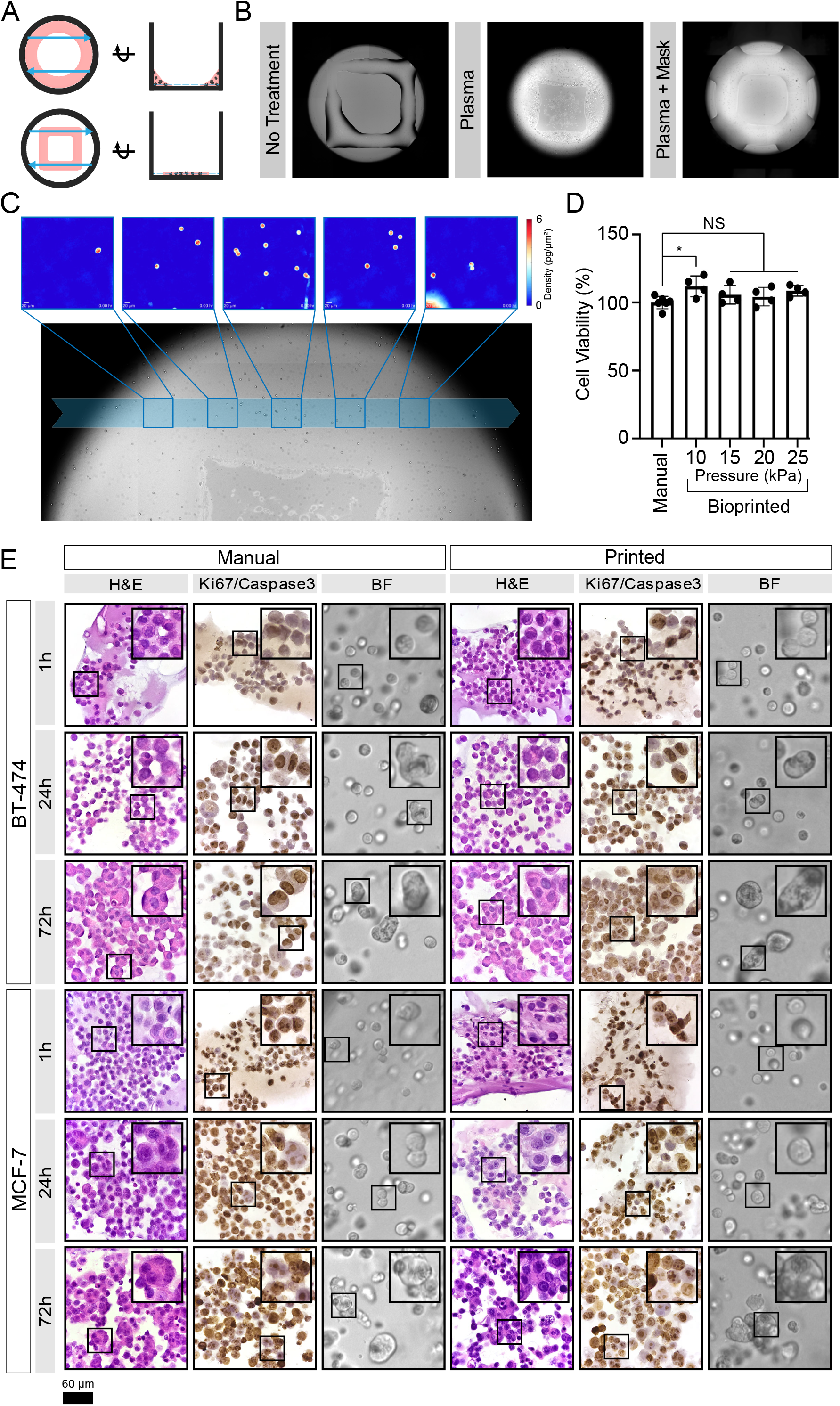
Bioprinting enables seeding of Matrigel-encapsulated organoids optimized for efficient HSLCI. (A) Schematic of wells with mini-rings (top) and mini-squares (bottom) relative to HSLCI imaging path (blue arrows). The top views (left) demonstrate that transitioning from rings to squares increases the area of material in the HSLCI imaging path. The side views (right) show that organoids in the square geometry align to a single focal plane better than organoids in a ring. (B) Plasma treatment of the well plate prior to printing optimizes hydrogel construct geometry. Bioprinting Matrigel onto untreated glass (left) generates thick (~200 *μ*m) constructs that decreases the efficiency of organoid tracking by increasing the number of organoids out of the focal plane. Whole well plasma treatment (middle) increases the hydrophilicity of all well surfaces causing the Matrigel to spread thin (~50 *μ*m) over the surface; however, the increased hydrophilicity also draws bioink up the walls of the well. Plasma treatment with a well mask facilitates the selective treatment of a desired region of the well (right). This leads to optimal constructs with a uniform thickness of approximately 75 *μ*m across the imaging path. (C) Individual organoids can be tracked over time across imaging modalities. Five representative HSLCI images are traced to the imaging path across a brightfield image. (D) Cell viability of printed versus manually seeded MCF-7 cells in a Matrigel-based bioink. A one-way ANOVA was performed (p = 0.0605) with post-hoc Bonferroni’s multiple comparisons test used to compare all bioprinted conditions against the manually seeded control. Adjusted p-values were 0.0253, 0.6087, >0.9999, 0.1499 for print pressures 10, 15, 20, 25 kPa, respectively. (E) H&E staining shows the development of multicellular organoids over time regardless of seeding method. The prevalence and size of multinuclear organoids increase with culture time. Ki-67/Caspase-3 staining demonstrates that most cells remain in a proliferative state throughout culture time. While apoptotic cells were observed in organoids cultured for 72 hours, the majority of cells show strong Ki-67 positivity. All images are 40X magnification and insets are 80X magnification. Ki-67 is stained brown, and caspase-3 is stained pink.

We next coupled these bioprinted organoids to an HSLCI platform. HSLCI uses a wavefront sensing camera and a dynamic focus stabilization system to perform continuous, high-throughput, label-free, quantitative phase imaging of biological samples, tracking their biomass changes over time^27,28^. However, efficient high-throughput QPI of 3D organoids using HSLCI is hindered by geometry considerations; when an object of interest is out of focus, measured phase shifts cannot be assumed to maintain a direct relationship with mass density^30^. Thus, we attempted to generate thinner layers of Matrigel to yield a relatively greater number of organoids in focus that can be quantitatively assessed at any given time. To generate thinner (<100 *μ*m) constructs amenable to efficient, label-free HSLCI imaging, we increased the hydrophilicity of the surface of 96-well glass-bottom plates by oxygen plasma treatment^41^. We developed 3D masks composed of BioMed Amber Resin (FormLabs) to selectively functionalize the region of interest (**Figure S1**). Bioprinting post-plasma treatment generated uniform mini-squares with organoids closely aligned on a single focal plane at ~70 *μ*m thickness (**Figure 1B**). These thin, printed mini-squares are amenable to massively parallel QPI by HSLCI as we aligned the legs of the bioprinted mini-square construct with the HSLCI imaging path (**Figure 1C**).

Lastly, we verified that the printing parameters used did not altered cell viability by directly comparing MCF-7 cells manually seeded according to our established protocol^6,7,21^ to cells printed through a 25G needle (260 *μ*m inner diameter) using extrusion pressures ranging from 10 to 25 kPa. We did not observe any reduction in cell viability as measured by ATP release assay (**Figure 1D**). These results are consistent with the existing literature as reductions in cell viability are often associated with higher print pressures (50-300 kPa)^42,43^. Taken together, this describes a method for bioprinting layers suitable for high-throughput HSLCI imaging without impacting cell viability, while supporting automated liquid-handling for high-throughput applications^6,7,21,44,45^.

### Bioprinted tumor organoids maintain histological features of manually seeded organoids

To verify that bioprinting did not perturb tumor biology, we directly compared the histology and immunohistochemical profiles of bioprinted and hand-seeded organoids from two breast cancer cell lines, BT-474 and MCF-7. These lines were selected for their differing molecular features such as their human epidermal growth factor receptor 2 (HER2) and estrogen receptor (ER) status^46^. We seeded cells as maxi-rings (1×10^5^ cells/ring) to obtain sufficient material for downstream characterization. Cells were either manually seeded into 24 well plates^6,7,21^ or bioprinted into 8-well plates at an extrusion pressure of 15 kPa. The bioprinted cells and resulting organoid structures were morphologically indistinguishable from manually seeded ones in brightfield images and hematoxylin and eosin (H&E)-stained sections taken 1, 24 and 72 hours after seeding (**Figure 1E**). Both bioprinted and manually seeded samples grew in size over time and bioprinting did not alter proliferation (Ki-67 staining) or apoptosis (cleaved caspase-3; **Figure 1E**). Hormone receptor status was unaltered, as shown by IHC for HER2 (**Figure S3**) and ER (**Figure S4**), and in agreement with literature reports for both cell types^47–50^. Thus, bioprinting did not influence organoid histology.

### Bioprinted and manually seeded organoids are molecularly indistinguishable

While bioprinted organoids are histologically indistinguishable from manually-seeded ones, this does not preclude molecular changes caused by the printing process. We therefore performed a detailed analysis of the transcriptomes of manually seeded and bioprinted cells 1-, 24- and 72-hours post-seeding. We assessed the distributions of 27,077 transcripts and clustered these into deciles based on their median abundance and found no significant differences between seeding approaches (**Figure 2A**). The overall transcriptomes of manually seeded and bioprinted organoids were extremely well-correlated (**Figure 2B**), with no individual transcripts differing significantly in abundance in either cell line even at very permissive statistical thresholds (0/27,077 genes, q <0.1, Mann-Whitney U-test, **Figure 2C**).

**Figure 2.**
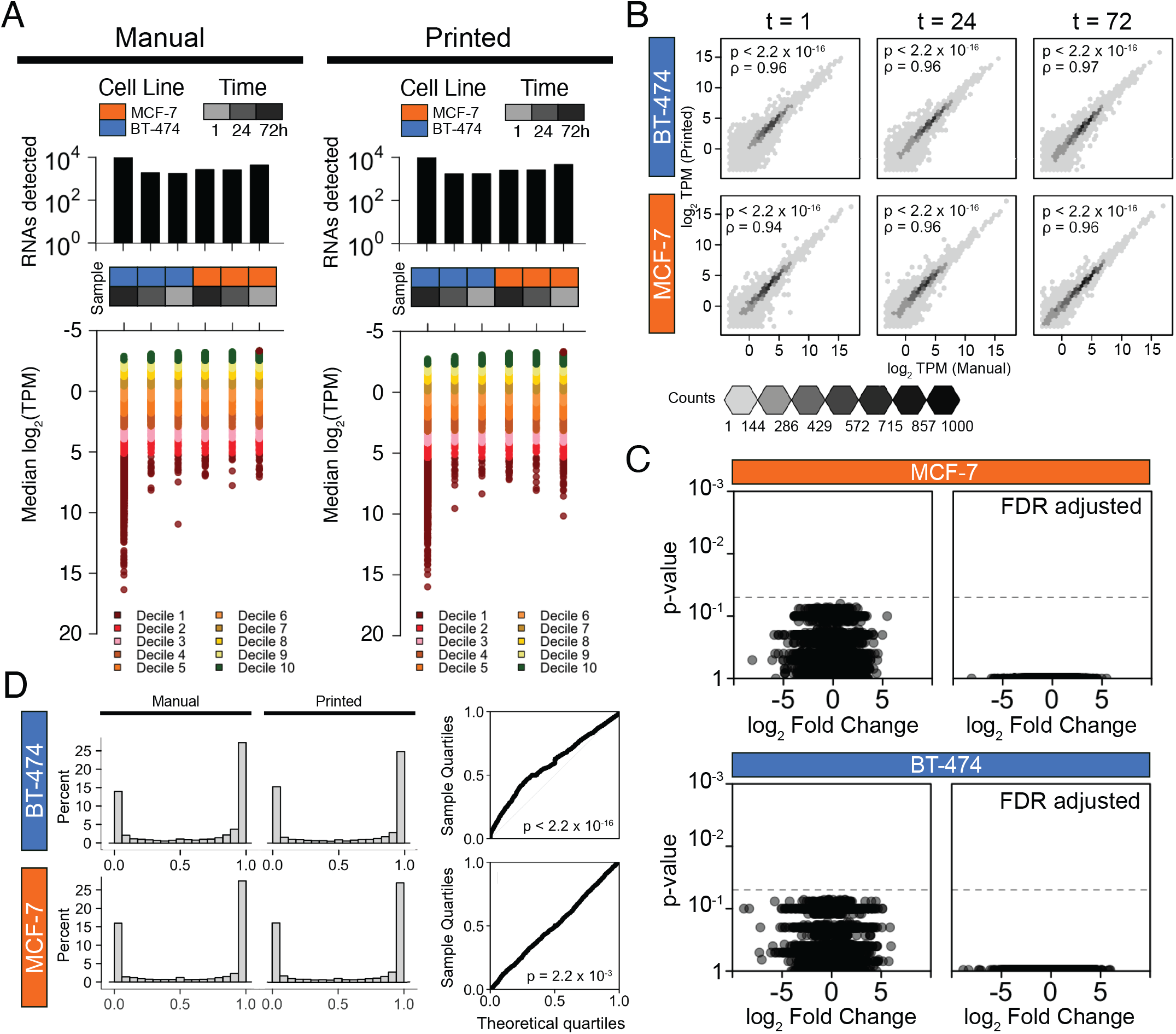
Bioprinting does not significantly alter organoid transcriptomes. (A) Distributions of total number of transcripts detected (above) and transcript abundances (below) measured as transcripts per million (TPM) organized into groups of deciles based on median abundance. (B) RNA abundances (log_2_ TPM) of manually seeded and bioprinted organoids at three different time points (t = 1, 24, and 72 hours). Spearman’s ρ was assessed for each association. We found strong associations between RNA abundances derived from printed and manually seeded organoids for both cell lines. (C) Volcano plots of Mann-Whitney U-test results for MCF-7 and BT-474 organoids with unadjusted p-values (left) and false discovery rate (FDR) adjusted p-values (right) comparing the RNA abundances of transcripts between manually seeded and printed tumor organoids. Fold change of RNA transcripts were assessed and log_2_ transformed. No transcripts were preferentially expressed based upon seeding method for organoids of either cell line (n = 0 out of 27,077 genes, q-value <0.1, Mann-Whitney U-test). (D) Median percent spliced in (PSI) of exon skipping isoforms were similarly distributed among BT-474 (top) and MCF-7 (bottom) derived organoids. Distribution of isoforms is consistent between manually seeded (left) and bioprinted (right) organoids. PSI of 1 indicates that the isoform is exclusively an exon inclusion isoform, while a PSI of 0 indicates that the isoform is exclusively an exon skipping isoform.

We next examined pre-mRNA alternative splicing events since these can induce functional changes even in the absence of variations in mRNA levels^51–53^. The density of exon-inclusion and exon-skipping isoforms was unchanged, with no individual fusion isoforms associated with the organoid printing method in either cell line (0/8,561, q <0.1, Mann-Whitney U-test; **Figure 2D**). Similarly, the number of fusion transcripts were not associated with seeding method (p = 0.17, Mann-Whitney U-test), although large numbers of fusions were detected in only one or two samples, reflecting the wide-spread *trans* splicing and genomic instability of immortalized cell lines^54^ (**Figure S5A**). Finally, there were no significant differences in the number or nature of RNA editing sites between printed and manually developed organoids (p = 0.48, Mann-Whitney U-test; **Figure S5B**). These findings demonstrate that our bioprinting protocol does not significantly impact the molecular characteristics of tumor organoids.

### Machine learning-based image segmentation and organoid classification enables single-organoid analysis

Our complete pipeline includes cell bioprinting (day 0), organoid establishment (day 0-3), full media replacement (day 3, **Figure 3A**) followed by transfer to the HSLCI incubator. Within 6 hours of media exchange, the plates are continuously imaged through 72 hours post-treatment. At the end of the imaging period, we perform an endpoint ATP assay to assess cell viability (**Figure 3A**). Interferograms collected by HSLCI are first converted to phase shift images using the SID4 software development kit (GPU version, v741)^30^. These images are then analyzed using two types of machine-learning algorithms.

**Figure 3.**
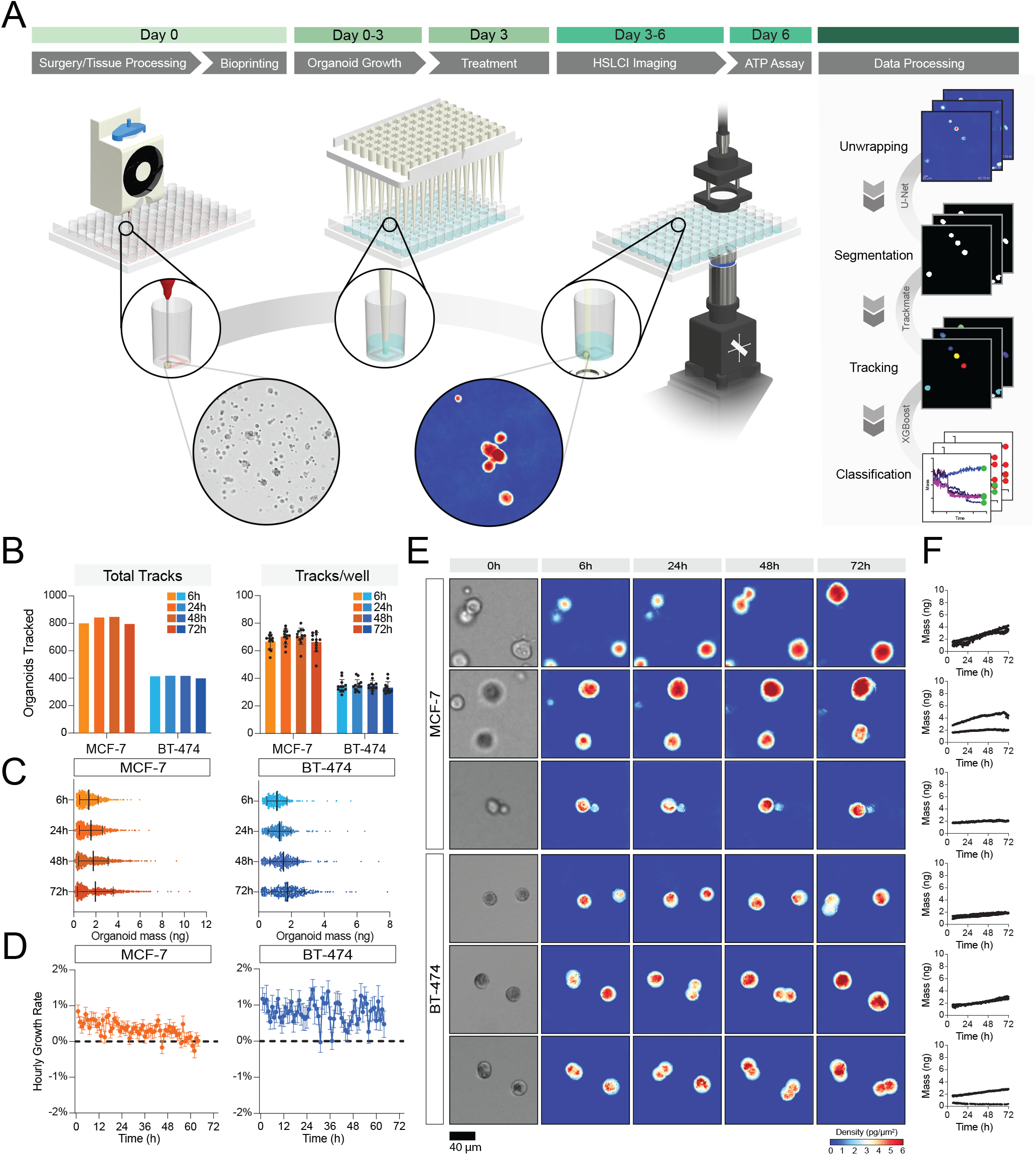
Bioprinting enables single-organoid tracking with high-speed live cell interferometry. (A) Extrusion-based bioprinting is used to deposit single-layer Matrigel constructs into a 96-well plate. Organoid growth can be monitored through brightfield imaging. After treatment, the well plate is transferred to the high-speed live cell interferometer for phase imaging. Coherent light illuminates the bioprinted construct and a phase image is obtained. Organoids are tracked up to three days using the HSLCI and changes in organoid mass are measured to observe response to treatment. (B) Total number of organoid tracks (left) and mean number of tracks per well (right) 6 hours (pale) and 48 hours (dark) after treatment for each cell line. The total number of organoid tracks across interpretable, replicate wells was 67 for MCF-7 organoids (n = 8), and 101 for BT-474 organoids (n = 12). At the 48-hour time-point the total number of tracks was 89 and 106 for MCF-7 (n = 8) and BT-474 (n = 10), respectively. (C) Mass distribution of tracked organoids 6 and 48 hours after treatment. Black bars represent the mean with error bars representing the standard deviation. (D) Hourly growth rate (percent mass change) of tracked MCF-7 (left) and BT-474 (right) organoids cultured in 1% DMSO. (E) Representative images of MCF-7 and BT-474 organoids tracked with HSLCI. Brightfield images of organoids taken immediately before treatment are shown on the left. (F) Calculated mass of each representative organoid over time.

To reliably identify unique organoids within each imaging frame despite the presence of background noise, debris, and out-of-focus organoids, we performed image segmentation using a U-Net architecture^55^ with a ResNet-34^7,56^ as the backbone. U-Net, a type of convolutional neural network (CNN), consists of an encoder that extracts rich feature maps from an input image and a decoder that expands the resolution of the feature maps back to the image’s original size. The long skip connections between the encoder and decoder propagate pixel-level contextual information into the segmented masks. The resulting segmentation images are very detailed even when provided small training datasets. The training dataset consisted of manually labeled organoids in 100 randomly selected imaging frames. This model created binary masks indicating whether each pixel of the image belonged to an organoid or the background with a mean Jaccard Index of 0.897 ± 0.109 at the 95% confidence level (**Figure S6**). The CNN reliably created masks omitting phase artifacts resulting from aberrant background or out-of-focus organoids (**Figure S6**).

Next, we determined the mass of each organoid in segmented masks by integrating the phase shift over the organoid area and multiplying by the experimentally determined specific refractive increment^29,34,36,37,57^. Organoids in subsequent frames were assembled into time coherent tracks using TrackMate^58,59^ and filtered using an XGBoost classifier^60^ we developed to exclude organoids moving in and out of focus, frequently overlapping and/or separating from neighboring organoids, or incorporating debris. We validated the model via cross-validation with 3-fold resampling of the sample population. The 3-fold resampling cross-validation score of the classifier was 91% with 93.5% accuracy. We also observed trends in the features used to classify each track, with excluded tracks typically have an increased number of missing frames as well as smaller interquartile ranges, and smaller initial and final sizes (**Figure S7**).

### Trends in mass accumulation of bioprinted organoids can be quantified by HSLCI with single-organoid resolution

HSLCI-based imaging allowed continuous tracking of n=921 MCF-7 organoids in 12 replicate wells (median: 78.5 organoids/well) and n=438 BT-474 organoids in 12 replicate wells (median: 36 organoids/well, **Figure 3B**). Due to organoids moving in and out of the field-of-view, the number of organoids tracked at each time point varied slightly. Overall, we tracked an average of 821 ± 28 MCF-7 organoids and 412 ± 9 BT-474 organoids at any given time throughout imaging (**Figure 3B**).

Unlike chemical endpoint assays or other live imaging modalities, HSLCI-based imaging facilitates parallel mass measurements of individual organoids. The initial average organoid mass was larger for MCF-7 (1.36 ± 0.84 ng) than BT-474 organoids (1.12 ± 0.61 ng, **Figure 3C**). The difference persisted throughout the entire imaging duration (**Table S1**). BT-474 cells grew at a rate of 0.80 ± 6.07% per hour while MCF-7 organoids demonstrated slower average hourly growth rates (0.33 ± 4.94% per hour, **Figure 3D**). The growth rate of the 3D BT-474 organoids is slightly slower than that observed after 6 hours in 2D culture (approximately 1.3%), while the MCF-7 organoids showed a much lower growth rate than previously reported 2D cultures (approximately 1.7%)^39^. We also observed positive associations between initial organoid mass and growth rate in both cell lines; however, the association between these factors is stronger for MCF-7 organoids (**Figure S8**). The varying degrees of association provide evidence of cell-line specific growth characteristics that cannot be measured using any other analytical method.

### Drug responses of organoids can be quantified by HSCLI

We then tested the utility of our platform in detecting drug responses in high-throughput 3D screenings (**Figure 3A**). As proof-of-principle we tested staurosporine, a non-selective protein kinase inhibitor with broad cytotoxicity^61^, neratinib, an irreversible tyrosine kinase inhibitor targeting EGFR and HER2^62^, and lapatinib, a reversible tyrosine kinase inhibitor also targeting EGFR and HER2^63^. Staurosporine and neratinib were tested at 0.1, 1, and 10 *μ*M, while lapatinib was screened at 0.1, 1, 10, and 50 *μ*M (**Figure 4 and S9**). These concentration ranges include and extend beyond the maximum plasma concentration reported for both lapatinib (4.2 *μ*M)^64^ and neratinib (0.15 *μ*M)^65^.

**Figure 4.**
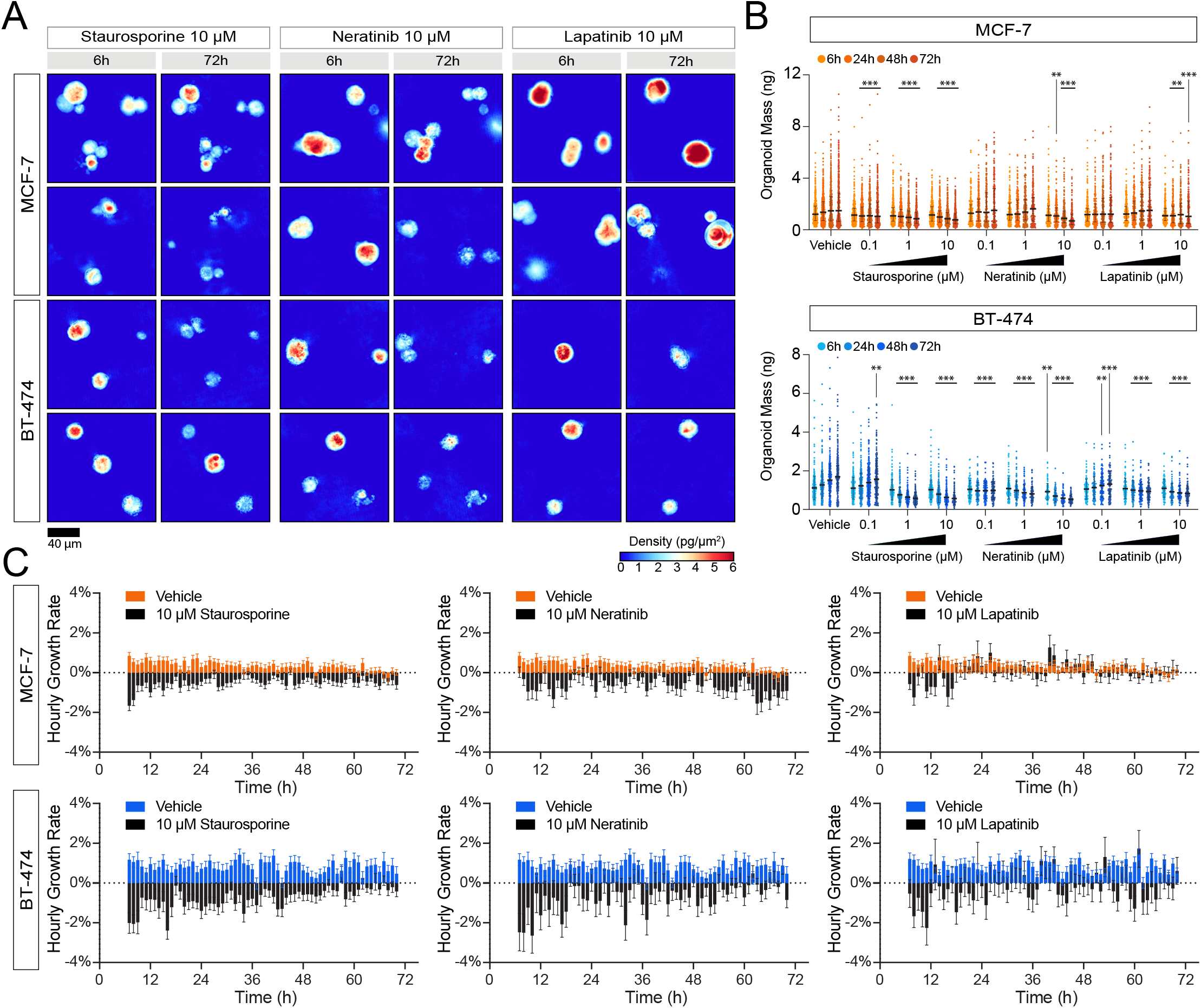
HSLCI enables high-throughput, longitudinal drug response profiling of 3D organoid models of cancer. (A) Representative images of organoids treated with 10 *μ*M staurosporine, 10 *μ*M neratinib, and 10 *μ*M lapatinib. (B) Mass of tracked MCF-7 and BT-474 organoids by treatment. Each bar represents the mass distribution at 6-, 24-, 28-, and 72-hours post-treatment (left to right). Black horizontal bars represent the median with error bars representing the interquartile range of the distribution. (C) Hourly growth rate comparisons (percent mass change) between organoids treated with 10 *μ*M staurosporine and vehicle, 10 *μ*M neratinib and vehicle, and 10 *μ*M lapatinib and vehicle. p<0.05 is denoted by *, p<0.01 is denoted by **, and p<0.001 is denoted by ***.

Representative HSLCI images demonstrate a range of responses to treatment (**Figure 4A**). The average masses at the start of the imaging window (6 hours post-treatment) did not significantly differ from the vehicle control (**Figure 4B, Table S1**). After 24, 48, and 72 hours, we observed significant differences in a number of treated samples (**Table S1**). After 24 hours, control MCF-7 organoids averaged 1.56 ± 1.05 ng, while those treated with 1 *μ*M and 10 *μ*M staurosporine showed significant reductions in average masses to 1.18 ± 0.77 ng (p = 1.93 x 10^-9^, Mann-Whitney U-test) and 1.11 ± 0.69 ng (p = 1.49 x 10^-13^, Mann-Whitney U-test), respectively. BT-474 organoids showed a similar pattern after 24 hours with control organoids averaging masses of 1.27 ± 0.69 ng while staurosporine-treated organoids averaged 0.76 ± 0.39 ng (1 *μ*M, p = 2.27 x 10^-32^, Mann-Whitney U-test) and 0.79 ± 0.44 ng (10 *μ*M, p = 1.59 x 10^-28^, Mann-Whitney U-test). The normalized growth curves (**Figure S10**) rapidly show response to treatment with 1 *μM* staurosporine.

Responses to lapatinib and neratinib reflected cell-specific trends. BT-474 organoids quickly showed sensitivity to both neratinib and lapatinib, while MCF-7 organoids only exhibited sensitivity to 10 *μ*M lapatinib and neratinib (**Figures 4B and 5A, Tables S1 and S2**). After 24 hours, the mean mass of the BT-474 organoids treated with 0.1 *μ*M neratinib decreased to 0.97 ± 0.44 ng from 1.27 ± 0.69 ng (p = 4.86 x 10^-6^, Mann-Whitney U-test) and organoids treated with 1 *μ*M lapatinib decreased to 1.00 ± 0.49 (p=3.37 x 10^-5^, Mann-Whitney U-test).

**Figure 5.**
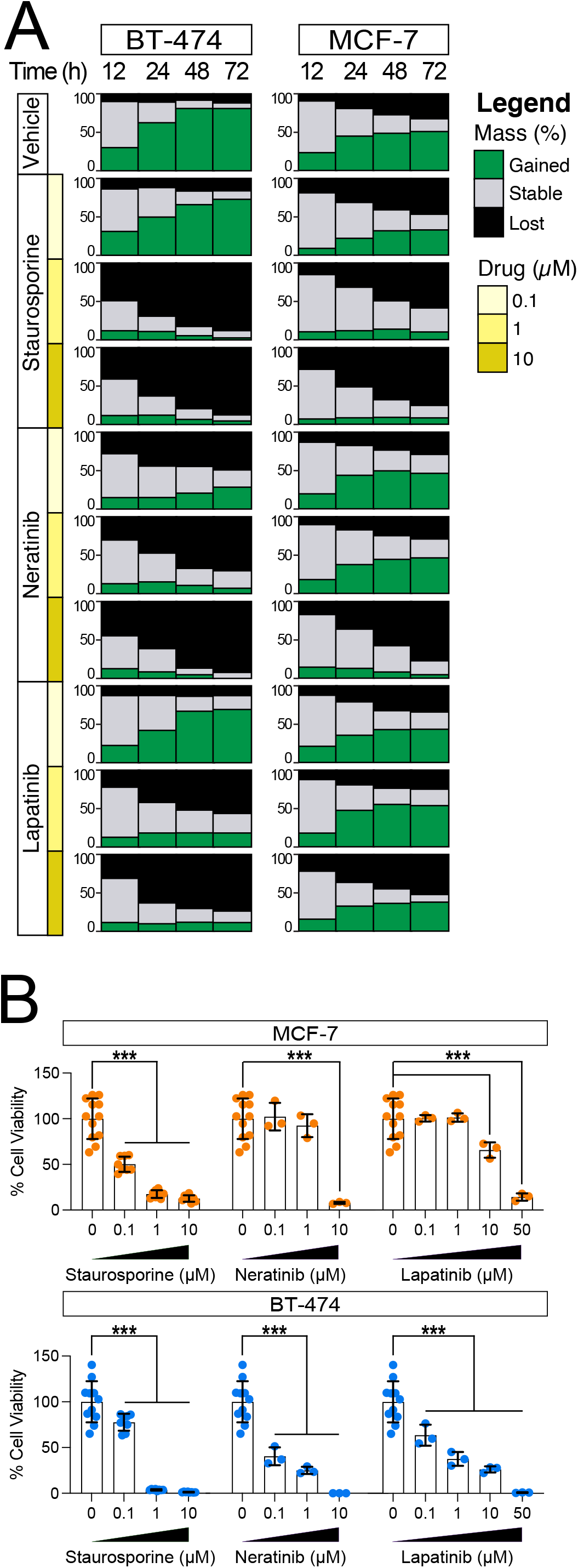
HSLCI enables identification of resistant and sensitive organoid subpopulations and discerns response to treatment earlier than a standard endpoint assay. (A) Plots showing the percentage of organoids in each condition that gain (green) or lose (black) more than 10% of their initial mass 12, 24, 48, 72 hours after treatment. (B) Percent cell viability of treated wells determined by an ATP-release assay. Statistical significance was assessed using an unpaired t-test with Welch’s correction. p<0.05 is denoted by *, p<0.01 is denoted by **, and p<0.001 is denoted by ***.

### Intra-sample heterogeneity of organoid drug responses

Our combination of HSLCI with ML-based organoid tracking provides per-organoid mass tracking, allowing quantitation of intra-sample heterogeneity (**Figure 3E-F, Supplementary Videos 1 & 2**). We assessed the ratio of organoids that gained, lost, and maintained mass over 12, 24, 48, and 72 hours for both control and treated samples (**Figure 5A, Table S2**). In the absence of drug treatment, 11.9% of BT-474 organoids lost more than 10% of their initial mass and 80.9% gained more than 10% of their initial mass over 72 hours. In contrast, only 50.8% of MCF-7 organoids gained mass and 32.6% lost mass. This heterogeneity in organoid populations increases over time, with 23.2% of MCF-7 organoids gaining more than 10% mass within 12 hours. This proportion nearly doubles to 44.8% after 24 hours but remains consistent at 48.6% and 50.8% after 48 and 72 hours, respectively. This pattern differs from BT-474 organoids as the population of organoids that gained mass continually increases over the first 48 hours before plateauing between 48 and 72 hours. BT-474 organoids that gained >10% mass increased from 30.1% after 12 hours, to 62.4% after 24 hours, and 80.9% after 48 and 72 hours.

Upon treatment, we could observe both inter-sample (MCF-7 vs BT-474) as well as intra-sample heterogeneity. In the presence of the HER2-targeting lapatinib (10 *μ*m), 37.7% of MCF-7 continued to grow and an additional 10.0% maintained their mass after 72 hours of treatment (**Figure 5A, Table S2**). When treated with 10 *μ*M neratinib, only 4.5% of MCF-7 organoids gained mass, while 18.1% remained stable. In contrast, BT-474 organoids showed greater sensitivity to both drugs, with 11.4% growing and 73.7% losing mass with 10 *μ*M lapatinib treatment (vs 11.9% for controls), and no organoids growing after 10 *μ*M of neratinib for 72 hours (**Figure 5A, Table S2**). A subset of BT-474 organoids showed high sensitivity to 0.1 *μ*M of both lapatinib and neratinib. In response to lapatinib, 12.8% of BT-474 organoids lost mass, while 17.9% maintained stable mass. When treated with neratinib, 49.1% lost mass and 22.4% had stable mass. Both responses contrasted with organoids treated with vehicle, of which 11.9% lost mass and 7.2% maintained mass. The heightened sensitivity of BT-474 cells to lapatinib and neratinib is expected given the higher expression of HER2 found in these cells^46^ (**Figure S3**).

A fraction of organoids in all treatment-cell combinations were unresponsive to the drugs tested (**Figure 5A, Figure S10, Table S2**). These organoids grew at similar rates to vehicle-treated cells and comprised between 7.8 and 87.1% of all organoids depending on the cell line and drug. For example, 37.7% of MCF-7 organoids treated with 10 *μ*M lapatinib grew after 72 hours, while an additional 10% maintained stable mass (**Table S2**). Similarly, when treated with 10 *μ*M neratinib for 72 hours, nearly 8% of BT-474 organoids maintained their mass, and when exposed to 10 *μ*M lapatinib for 72 hours, the proportion increases to over 25% (**Table S2**). Our findings are indicative of a resistant population of organoids that can be rapidly identified by HSLCI imaging. These persisters may provide a unique model for understanding *de novo* and acquired treatment resistance.

Lastly, to validate the responses measured by HSLCI, we performed an endpoint ATP-release assay on the same plates used for HSLCI imaging and assessed organoid viability at the end of the 72-hour treatment (**Figure 5B**). The ATP assay confirmed that both cell lines are highly sensitive to staurosporine with near-zero viability at the 1 and 10 *μ*M concentrations. Additionally, BT-474 organoids show significant reductions in viability when treated with 0.1 *μ*M lapatinib and 0.1 *μ*M neratinib for 72 hours (**Table S3**). Overall, the results of the cell viability assay after 72 hours confirm the trends observed in as little as 6 hours by HSLCI but fail to capture intra-sample variability.

## Discussion

Every newly diagnosed human cancer reflects a unique set of germline variation, somatic mutations, and microenvironmental influences^66^. Cancer therapy attempts to address this by personalizing treatment for individual patients^67,68^. The most common approach to date has been molecular precision medicine, which links therapeutic efficacy to molecular features of a tumor^1,69,70^. Functional precision medicine approaches, by contrast, bypass the need to learn drug-molecular associations by relating *ex vivo* response to clinical outcomes^8,71,72^. Key limitations towards the broad adoption of functional precision medicine have been the creation of physiological culture models, the development of high-throughput systems, and the difficulty in measuring organoid heterogeneity^3,73,74^. Here, we describe a new pipeline that overcomes these barriers by incorporating a robust 3D organoid bioprinting protocol and an imaging approach that facilitates single-organoid analysis of response to treatment.

We introduced bioprinting to enhance the throughput and consistency of our previously published organoid screening approaches^6,7,21^. We opted to print a Matrigel-based bioink due to its ability to preserve tumor characteristics *ex vivo^6^;* however, its weak mechanical integrity and its temperature-dependent viscosity and crosslinking behavior complicate its suitability for bioprinting^75^. To circumvent these limitations, we optimized a protocol that takes advantage of its temperature-dependent behavior to yield consistent mechanical properties for bioprinting. While the existing consensus is that consistent bioprinting with Matrigel is difficult to achieve, we show that simple, single-layer structures are attainable with strict temperature regulation. We further enhanced the quality of the Matrigel deposition by selectively modifying the print substrate with oxygen plasma treatment. The introduction of 3D plasma masks (**Figure S1**) facilitated the selective treatment of a square region in each well. The increased hydrophilicity of the substrate in the exposed region guides the spreading of the material to ensure maximize consistency in deposition volume and construct thickness while preventing obstruction of the center of the well. Bioprinting allowed us to finely control the size and shape of the deposited gel constructs, facilitating the use of HSLCI for downstream analysis (**Figure 1**).

To our knowledge, this is the first reported use of live cell interferometry for label-free, time-resolved quantitative imaging of 3D organoid cultures. Previous studies have used interferometry to quantify the mass of individuals cells cultured on 2D substrates to study cell division^76^, cytoskeletal remodeling^77^, mechanical properties^78^, and response to treatment^27–29^. Tomographic QPI has also been used to obtain high-resolution images of 3D objects such as cerebral organoids^79^. The primary challenge of adapting live cell interferometry for the mass quantitation of 3D organoids is maintaining the organoids in a single focal plane. The mass of organoids outside of the focal plane cannot be accurately calculated as phase information for out-of-focus planes is difficult to interpret^30^. We were able to circumvent this challenge by introducing bioprinting to generate uniform, thin constructs that maximize the number of organoids that could be tracked in parallel, and a machine-learning based organoid classifier to exclude out-of-focus and non-organoid objects from our analysis. By introducing region-specific reference images and machine-learning based methods for image segmentation and track filtering, we have been able to increase the number of organoids tracked approximately 15-fold from initial analyses where only approximately 1% of organoids analyzed were retained per well using standard approaches. Further improvements will include shortening the 6-hour delay between drug treatment and imaging start, which will allow us to capture highly sensitive organoids undergoing cell death within that timeframe. Lastly, due to the large amount of data generated using HSLCI (approximately 250 GB per plate/day), data analysis remains a time-limiting factor.

Despite the development of 3D cancer models with varying extents of complexity and scalability, functional screening assays have been hindered by their inability to consider the heterogeneity of tumor response. Genomic characterization of tumors has demonstrated that these malignancies are collections of evolutionarily-related subclones, rather than homogeneous populations^80–83^. This genetic diversity is one of the several factors that contributes to differential response to treatment. Endpoint assays, such as live-deadstaining or ATP-release quantification, characterize the average response to treatment. Though they may be useful for identifying drug sensitivity in majority cell populations, they fail to account for the response of resistant populations that may also be present. In the clinical setting, failure to treat the resistant populations may lead to initial response, followed by recurrence and long-term disease progression^84–86^. HSLCI allows us to non-invasively track various features of the bioprinted organoids over time, including size, motility, and mass density. Because of the ability to quantitatively measure mass changes in response to treatment, it is possible to identify and isolate responsive and resistant subpopulations of cells, which can in turn lead to better informed clinical decision making.

## Acknowledgments

We acknowledge the UCLA Translational Pathology Core Laboratory and the UCLA Technology Center for Genomics and Bioinformatics for their assistance with this work. Additionally, we would like to thank Dr. Steven Jonas for his generous access to laboratory equipment. This work was supported by a NIH R01CA244729 (to A.S. and P.C.B.), a National Science Foundation Graduate Research Fellowship (DGE-2034835, to B.W.), a Eugene V. Cota Robles Fellowship (to B.W.) a UCLA DGSOM Seed Award (to A.S. and P.C.B.), the Air Force Office of Scientific Research (FA9550-15-1-0406, to M.A.T.), the Department of Defense (W81XWH2110139, to M.A.T.). It was additionally supported by NIH U24CA248265 (to P.C.B.) and NIH R01GM114188, R01GM073981, R01CA185189, R21CA227480, R01GM127985, and P30CA016042 (to M.A.T.). A.S., P.J.T., B.W. and N.T. are inventors on a patent application based on some aspects of this work. P.C.B and A.S. are founders and owners of Icona BioDx.

## Materials and Methods

### 2D Cell Culture

MCF-7 and BT-474 breast adenocarcinoma cell lines were obtained from the American Type Culture Collection (ATCC). All cell lines were grown for a maximum of 10 passages in RPMI 1640 (Gibco 22400-089) supplemented with 10% fetal bovine serum (FBS, Gibco 16140-071) and 1% antibiotic-antimycotic (Gibco 15240-062). Both cell lines were periodically authenticated by short tandem repeat profiling using the GenePrint 10 kit (Laragen).

### Manually Seeded 3D Organoids

Organoids were seeded manually according to our previously published protocols^6,7,21^. Briefly, single cells suspended in a 3:4 mixture of Mammocult (StemCell Technologies 05620) and Matrigel (Corning 354234) were deposited around the perimeter of the wells of either 24-well or 96-well plates. The cell suspension was kept on ice throughout the seeding process to prevent gelation of the Matrigel. To seed organoids in a 96-well plate (Corning 3603), a pipette was used to distribute 5 *μ*L of cell suspension (5×10^5^ cells/mL) along the bottom perimeter of each well. Once all mini-rings are generated, plates were incubated at 37°C and 5% CO_2_ for 20 minutes to solidify the Matrigel, and 100 *μ*L of pre-warmed Mammocult was added to the center of each well using an epMotion 96 liquid handler (Eppendorf). To generate larger rings (maxi-rings) in 24-well plates (Corning 3527), 70 *μ*L of cell suspension (1.4×10^6^ cells/mL) was deposited around the perimeter of each well. Following seeding, the plate was incubated at 37°C and 5% CO_2_ for 45 minutes to solidify the Matrigel, and 1mL of pre-warmed Mammocult was added to the center of each well.

### 3D Printing Plasma Masks

Custom well masks were designed to meet the specifications of the well plates that were used in these experiments (**Figure S1**). The design was generated in Inventor 2020 (Autodesk) and printed using a Form3B (FormLabs) using the Biomed Amber resin (FormLabs). The design was exported as an STL file and imported into the PreForm (FormLabs) software to arrange the parts. After printing, parts were post-processed in two washes of isopropanol, air-dried for at least 30 minutes, and cured for an additional 30 minutes at 70°C in the Form Cure (FormLabs).

### Bioprinted 3D Organoids

Cells were bioprinted using a CELLINK BioX with a Temperature-Controlled Printhead. Gcode files were written to print the desired single-layer geometry. MATLAB (MathWorks, Inc.) was used to integrate these standardized blocks into full Gcode files with the defined coordinates for each well. We used 8-well plates when printing the maxi-rings for IHC and RNA sequencing (RNAseq) as the depth of the well in a standard 24-well plate prohibited the use of 0.5” length needles. Four rings with a diameter of 14.5mm were printed for RNAseq (~2×10^5^ cells total), while four sets of concentric 14.5mm, 12.5mm, and 10.5mm diameter rings were used for IHC analysis (~5×10^5^ cells total). We printed mini-squares with side length 3.9mm for drug screening and HSLCI imaging. The mini-squares were inscribed within the circular well with sides parallel to the sides of the well plate. All bioprinting processes utilized the same material deposited for manually seeded organoids: a single-cell suspension in a 3:4 mixture of Mammocult and Matrigel on ice. After vortexing briefly, the mixture was transferred into a 3 mL syringe to remove air bubbles. The mixture was then transferred to a room temperature 3 mL bioprinter cartridge (CELLINK) by connecting the syringe and cartridge with a double-sided female Luer lock adapter (CELLINK). The loaded cartridge was incubated in a rotating incubator (Enviro-Genie, Scientific Industries) for 30 minutes at the print temperature.

During the incubation period, the printer was sterilized with the built-in UV irradiation function, the printhead was set to the print temperature and the masked 96-well plates treated with oxygen plasma. Briefly, well masks were autoclaved prior to use, inserted into the well plate, and pressed in contact with the glass surface. Masked plates were treated with oxygen plasma in a PE-25 (Plasma Etch) for 30-90 seconds, 15 minutes prior to bioprinting. After plasma treatment, the well plate was placed in the bioprinter and Automatic Bed Levelling (ABL) was performed.

Once the incubation period ended, we attached a 0.5” 25-gauge needle and loaded the cartridge into the pre-cooled printhead. We primed the needle by extruding a small volume of material at 15 kPa prior to calibrating the printer. The material in the needle gelled during the printer calibration which takes approximately 2 minutes. After calibration, we performed a second extrusion using 40 kPa to clear the needle of the gelled material prior to starting the print, this step ensured that we achieved unobstructed material extrusion. To create constructs of the appropriate thicknesses, prints in 8-well plates were extruded at 15 kPa while prints in 96-well plates were extruded at 12-15 kPa. The bioprinter completes the deposition process for 96-well plates in approximately four minutes. After printing, the constructs were incubated at 37°C for at least 30 minutes to solidify the matrix and 100μL of Mammocult medium was then added.

### Sample Preparation for RNA Sequencing

Organoids were released from the Matrigel in preparation for RNAseq. After aspirating the media from each ring, 1 mL of cold Dispase was added per ring. After a 20-minute incubation at 37°C, the cell suspension was collected and pelleted by centrifugation at 1500g for 5 minutes and washed with 45 mL of PBS before centrifuging again at 2000g for an additional 5 minutes. Once all liquid was aspirated, the tubes were rapidly frozen and stored at −80°C. Frozen cell pellets (~2×10^5^ cells) were then transferred to the Technology Center for Genomics & Bioinformatics (TCGB) at UCLA for RNAseq. Sequencing was performed on a NovaSeq SP (Illumina) using the 2 x 150 bp paired-end protocol.

### RNA Sequencing Data Processing and Analysis

FASTQ files were processed using UCLA-CDS pipelines to align, quantify and call RNA-sequencing reads. Pipeline-align-RNA v6.2.2 aligns paired-end, reverse stranded RNA-seq reads using STAR v2.7.6^87^ and HISTA2 v2.2.1^88^. Genome reference file, CRCh38. p13, was used for aligners STAR and HISTA2. Annotations were performed using Gencode v34 reference GTF. FASTP v0.21.0^89^ was included in the pipeline to trim reads for low-quality bases and remove adaptor sequences. Next, the pipeline marked duplicate reads using the GATK Spark tools^90^ (MarkDuplicates Spark v4.1.4.1^90^) that allowed for parallel processing on multiple computing clusters. Lastly, the pipeline runs dupRader v1.24.0^91^ to check the duplication rate.

We used pipeline-quantitate-RNA to quantify RNA at the gene and transcript isoform level. The pipeline used RSEM v1.3.3^92^ to quantify RNA using GRCh38.p13 as the reference index file. RSEM quantifies aligned RNA-seq in BAM format. The output is a quantitated RNA at the RNA and transcript isoform level. Quality control procedures include running FastQC v0.11.9 on input FASTQ files to control for low quality reads that may lead to low quality mapping. Transcripts with low abundance in all samples (TPM < 0.1; transcripts per million) were excluded resulting in 27,077/67,060 transcripts included in the analysis.

Pipeline-quantitate-SpliceIsoforms was used to quantitate the relative usage of splice isoforms using aligned RNA-seq data. The pipeline validates inputs and used rMATS v4.1.0^93^ on individual RNA-seq aligned data in BAM format. The output includes information on the alternative splicing event types. We excluded splice isoforms with missing data in five or more samples (8,561/17,449) due to low power.

Pipeline-call-RNAEditingSite uses REDItools2 v1.0.0^94^ to call RNA editing events. Pipeline-call-FusionTranscripts calls gene fusion events using a combination of Arriba v2.1.0^95^, STAR-Fusion v1.9.1^96^ and fusioncatcher v1.33^97^. Arriba detects gene fusions from RNA-seq data using the STAR aligner. The STAR-Fusion caller is a component of the Trinity Cancer Transcriptome Analysis Toolkit (CTAT). Fusioncatcher calls somatic fusion genes in paired-end RNA-seq data files. RNA editing sites were filtered to include adenosine to inosine events with sufficient coverage (q30 >10) and frequencies above 0.9. Poly-A depleted RNA included annotated microRNAs (miRNA), while poly-A enriched RNA included coding mRNAs. Raw and processed data will be made available in GEO.

We used a Mann-Whitney U-test to perform non-parametric hypothesis statistical testing of RNA abundances, number of transcript fusions, and editing sites between bioprinted and manually seeded tumor organoids. We adjusted for multiple hypothesis testing using the false discovery rate (FDR) method, setting q < 0.1 as the criterion for strong associations. Statistical analyses and data visualization were performed in the R statistical environment (v4.0.2) using the BPG^98^ (v6.0.1) package.

### Immunohistochemistry

Immunohistochemical staining was performed on manually seeded and bioprinted organoids seeded in 24 or 8-well plates, respectively. A detailed procedure has been published^21^. Briefly, samples were prepared for histological analysis by carefully aspirating all media from the well without disrupting the construct and fixing in 10% buffered formalin (VWR 89370-094). The fixed organoids were harvested, transferred to a conical tube and pelleted by centrifugation at 2000xg for 5 minutes. HistoGel (Thermo Scientific HG-40000-012) was then added to the pellet. Once solidified, the cell pellet in HistoGel was placed in a histologic cassette and sent to the UCLA Translational Pathology Core Laboratory (TPCL) for dehydration and paraffin embedding.

Slides (8 *μ*m thin sections) were baked for 20 minutes at 45 °C and de-paraffinized in xylene followed by washes in ethanol and deionized water. For H&E staining, a Hematoxylin and Eosin Stain Kit (Vector Labs H-3502) was used according to the manufacturer’s protocol. For Ki-67/Caspase-3, HER2, and ER staining, Peroxidazed-1 (Biocare Medical PX968M) was applied for 5 minutes at room temperature to block endogenous peroxidases. Next, antigen retrieval was performed using Diva Decloaker (Biocare Medical DV2004LX) in a 2100 Retriever (Prestige Medical) heating at 110 °C for 15 minutes. Blocking was performed at room temperature for 5 minutes with Background Punisher (Biocare Medical BP947H), Primary Ki-67/Caspase-3 staining was performed overnight with pre-diluted Ki-67/Caspase-3 (Biocare Medical PPM240DSAA) solution at 4°C after an additional 2-minute Background Punisher treatment post-antigen retrieval, and secondary staining was performed with Mach 2 Double Stain 2 (Biocare) solution for 40 minutes at room temperature. Primary antibodies for HER2 (Novus Biologicals, CL0269) and ER (Abcam, E115) staining were diluted 1:100 in Da Vinci Green Diluent (Biocare Medical PD900L). The HER2 antibody was incubated overnight at 4°C while the ER antibody was incubated at room temperature for 30 minutes. Secondary staining was performed with Mach 3 Mouse Probe and Mach 3 Mouse HRP-Polymer for HER2 and Mach 3 Rabbit Probe and Mach 3 Rabbit HRP-Polymer for ER (10 minutes). Chromogen development was performed with Betazoid DAB (Biocare Medical, BDB2004) followed by counterstaining with 20% hematoxylin (Thermo Scientific #7221). Slides were dehydrated in ethanol and xylene and coverslipped with Permount (Fisher Scientific SP15-100). Imaging was performed with a Revolve microscope (Echo Laboratories). Whole image white balancing was performed in Adobe Photoshop.

### Drug Screening

A detailed protocol for the drug screening has been published previously^6,21^. Briefly, the culture medium was fully removed three days after seeding and replaced with 100 *μ*L of Mammocult medium containing the indicated drug treatments using a liquid handler (EpMotion® 96). After treatment, we transferred the plates to the HSLCI platform for imaging.

### High-Speed Live Cell Interferometry

HSLCI has been described previously^27,28^. The HSLCI platform is a custom-built inverted optical microscope coupled to an off-axis quadriwave lateral shearing interferometry (QWLSI) camera (SID4BIO, Phasics, Inc.)^30^. This wavefront sensing camera incorporates a modified Hartmann mask that splits the incident wave front into four tilted replica wavefronts that interfere with one another. The resulting interferograms are recorded and used to recover phase gradients along two perpendicular directions, allowing for reconstruction of a phase shift map and subsequent calculation of dry mass of discrete objects within imaging fields of view (FOVs)^30,33^. Illumination is provided by a 660 nm fiber-coupled LED (Thorlabs). The HSLCI platform captures images from standard-footprint (128×85 mm) glass-bottom multiwell plates. Motorized stages (Thorlabs) control the xy-motion of a single glass-bottom plate above the microscope objective, and in combination with a piezo-actuated dynamic focus stabilization system, enable continuous and repeated image collection over many FOVs within each row of wells. The HSLCI platform is installed inside of a standard cell culture incubator to enable long-term imaging of samples in physiology-approximating conditions (37°C, 5% CO2). All hardware and software components are available commercially.

For all growth kinetics and drug screening studies, organoids were imaged in 96-well glass-bottom plates (Cellvis P96-1.5H-N) using a 40× objective (Nikon, NA 0.75). Plates were prepared as described and wrapped with parafilm to limit evaporation during imaging. Organoids were imaged continuously from 6 hours to 72 hours following administration of drug treatments. During imaging, the sample plate was translated along each row of wells such that about 25 images per well were collected on each imaging loop, and that imaging FOVs overlapped with areas of wells in which bioprinted matrix and organoids were present. The typical imaging interval was 10 minutes between successive frames at the same FOV.

### Machine Learning-Based Analysis of HSLCI Images

Images were acquired using the SID4Bio software (v2.4.2.93, Phasics). After image collection, interferograms captured by the QWLSI camera were converted to phase shift images using the SID4 software development kit for MATLAB (GPU version, v741, Phasics), in a process called phase unwrapping. For every frame at each FOV, phase shift maps were processed by converting to optical path difference^30^, and then subtracting the fourth-order Zernike polynomial^99^ fit to each frame using least-squares over a cartesian grid to remove refractive aberrations.

The processed images were then segmented into individual cells or organoids using a convolutional neural network (U-Net architecture^55^ with a ResNet-34 encoder^7,56^). We initialized our model with weights derived from a model pretrained on the ImageNet dataset^100^. The training dataset consisted of a randomly selected set of 50 images from the BT-474 dataset, and 50 from the MCF-7 dataset encompassing images taken at all wells, intra-well imaging positions, and timepoints within each experiment. Each 514×514 pixel image was overlaid with a binary mask that marks each pixel as either part of an organoid or the background. Using the training dataset, the weights were refined using a cross-entropy loss function over 80 epochs. We organized phase images and their corresponding masks into one stack per imaging FOV and sorted each layer by time of imaging. We used TrackMate (v7.6.1)^58,59^ to track the organoids over time using a Sparse LAP Tracker with a maximum linking distance of 90 pixels and feature penalties of 4.0 for both the major ellipse axis and organoid area. These parameters were selected based on optimal performance on a set of 12 representative image stacks. We tolerated gaps of up to 30 frames to maximize track continuity between organoids. Tracks composed of fewer than 10 frames were excluded. Labeled image stacks were exported and mass was extracted from the segmented regions by integrating the phase shift over the area and multiplying by the refractive increment of 1.8×10^-3^ m^3^/kg ^29,34,36,37,57^.

We developed an XGBoost-based classifier (v1.5.2.1, probability prediction type, log-loss evaluation metric)^60^ to predict permissibility of organoid tracks using R package mlr3 (v0.13.2). For the supervised learning model, we labelled a subset of tracks for permissibility (n = 846) from four wells: 1) BT-474 well treated with vehicle, 2) BT-474 well treated with 10 *μ*M staurosporine, 3) MCF-7 well treated with vehicle, and 4) MCF-7 well treated with 10 *μ*M staurosporine and manually determined that n = 250/846 tracks were acceptable for downstream analysis. We extracted a set of time-series based features from the mass reconstruction data to use in our classifier model. The features were number of missing frames, initial size, interquartile range (IQR), IQR of the first 12 points, and IQR of the last 12 points. Initial size was calculated based on the median mass of the first two timepoints (**Figure S2**). Area under the curve (AUC) measurements were calculated using R package (bayestestR v0.11.5). Pair-wise correlation plots comparing the void and valid track features were generated using mlr3viz R package (v0.5.7). We used probability as the learner prediction type and log-loss (log_10_) as the evaluation metric. To validate the model, we performed k-fold cross-validation protocols with 3-fold resampling. The classifier predicted n = 8,590/29, 137 to be permissible organoid tracks for downstream analysis with an accuracy of 93.5% and a cross-validation score of 91%.

### ATP release assay

Manually seeded organoids were prepared in accordance with the protocol described above and published^6,7,21^. To assess the viability of bioprinted organoids, we prepared the bioink and bioprinter as described. We extruded 100 *μ*L of bioink into and Eppendorf tube for each print pressure (10, 15, 20, and 25 kPa). We seeded four 10 *μ*L rings in a 96-well plate using the extruded bioink. For drug screenings, plates were retrieved from the HSLCI incubator and processed as briefly described. After a PBS wash, 50 *μ*L of 5 mg/ mL Dispase (Life Technologies 17105-041) solution was added to each well and incubated for 25 minutes. After shaking for 5 minutes on an orbital shaker at 800 RPM, we added 75 *μ*L of CellTiter-Glo® Luminescent Cell Viability Reagent (Promega G968B) to each well and followed the manufacturer’s instructions. Luminescence was measured using a SpectraMax iD3 (Molecular Devices) plate reader (parameters: read all wavelengths, signal integration of 500ms). The viability of each well was calculated by normalizing the luminescent signal to the average signal from the manually seeded control wells. An unpaired t-test with Welch’s correction was performed in GraphPad Prism. P-values less than 0.05 were deemed significant.

## Supplementary Material

### Supplementary Videos

**Supplementary Video 1.**
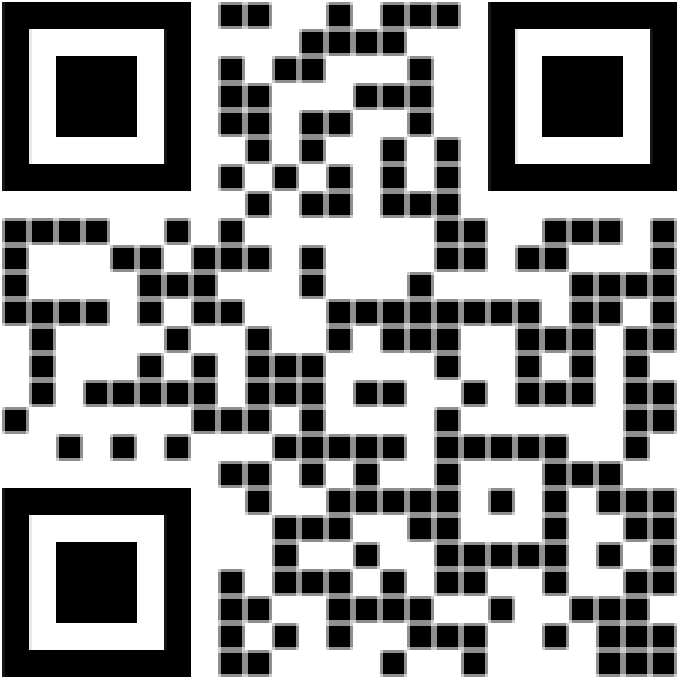
MCF-7 organoids treated with the vehicle control. Scan the QR code to access the video or visualize at the following link: https://youtu.be/bUBq-ZChFM0

**Supplementary Video 2.**
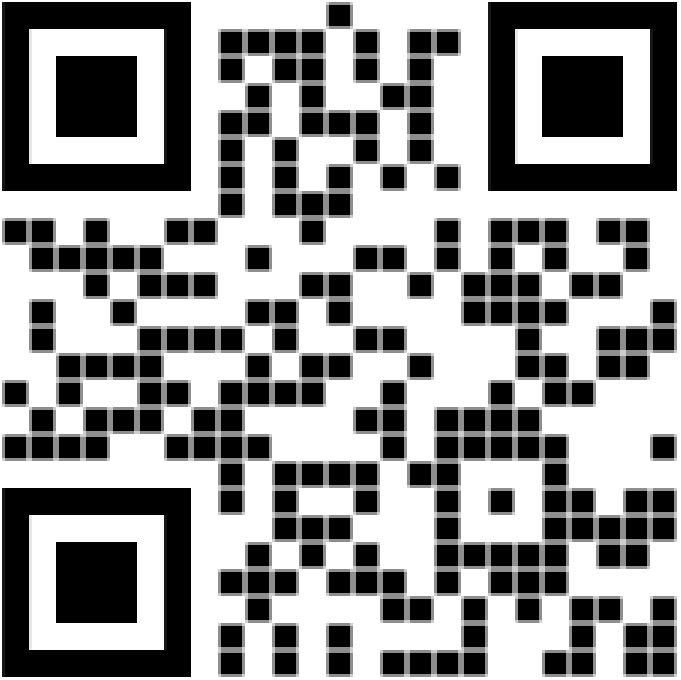
BT-474 organoids treated with the vehicle control. Scan the QR code to access the video or visualize at the following link: https://youtu.be/AzSc8WW5KBA

### Supplementary Tables

**Table S1:** Organoid mass distributions. (A) Comparisons of mean mass of MCF-7 organoids calculated via HSLCI. We first performed a Kruskal-Wallis test to determine if one sample stochastically dominates another. If the p-value was less than 0.05, we then performed Mann-Whitney U-tests for each sample against the vehicle control at the respective time point. Data is presented in Figure 6B. (B) Comparisons of mean mass of BT-474 organoids calculated via HSLCI. We first performed a Kruskal-Wallis test to determine if one sample stochastically dominates another. If the p-value was less than 0.05, we then performed Mann-Whitney U-tests for each sample against the vehicle control at the respective time point. Data is presented in Figure 3C.

|  |  | 6 hours | 24 hours |  | 48 hours |  | 72 hours |  |
| --- | --- | --- | --- | --- | --- | --- | --- | --- |
| MCF-7 | | Mean mass ± SD | Mean mass ± SD | p-value | Mean mass ± SD | p-value | Mean mass ± SD | p-value |
| Vehicle | 1% DMSO | 1.36 ± 0.84 | 1.56 ± 1.05 | - | 1.77 ± 1.33 | - | 1.95 ± 1.61 | - |
| Staurosporine | 0.1 μM | 1.30 ± 0.84 | 1.27 ± 0.89 | 2.11 $\times 10^{-5}$ | 1.39 ± 1.07 | 1.31 $\times 10^{-5}$ | 1.44 ± 1.21 | 2.49 $\times 10^{-6}$ |
| | 1 μM | 1.28 ± 0.84 | 1.18 ± 0.77 | 1.93 $\times 10^{-9}$ | 1.10 ± 0.73 | 2.24 $\times 10^{-17}$ | 1.03 ± 0.70 | 2.80 $\times 10^{-21}$ |
| | 10 μM | 1.30 ± 0.78 | 1.11 ± 0.69 | 1.49 $\times 10^{-13}$ | 1.01 ± 0.65 | 6.73 $\times 10^{-24}$ | 0.93 ± 0.64 | 3.24 $\times 10^{-30}$ |
| Neratinib | 0.1 μM | 1.47 ± 1.03 | 1.62 ± 1.18 | 1 | 1.86 ± 1.51 | 1 | 2.03 ± 1.74 | 1 |
| | 1 μM | 1.45 ± 0.96 | 1.57 ± 1.09 | 1 | 1.68 ± 1.22 | 1 | 1.79 ± 1.40 | 1 |
| | 10 μM | 1.39 ± 1.07 | 1.30 ± 0.98 | 0.0036 | 1.16 ± 0.92 | 2.90 $\times 10^{-9}$ | 0.92 ± 0.77 | 6.82 $\times 10^{-19}$ |
| Lapatinib | 0.1 μM | 1.38 ± 1.03 | 1.49 ± 1.19 | 0.9072 | 1.73 ± 1.47 | 1 | 1.79 ± 1.56 | 1 |
| | 1 μM | 1.40 ± 0.83 | 1.61 ± 1.11 | 1 | 1.84 ± 1.38 | 1 | 2.02 ± 1.69 | 1 |
| | 10 μM | 1.29 ± 0.83 | 1.30 ± 0.98 | 0.0011 | 1.42 ± 1.11 | 0.0013 | 1.50 ± 1.29 | 0.0006 |
| Kruskal-Wallis p-value | | 0.3169 | 2.21 $\times 10^{-22}$ | | 4.57 $\times 10^{-45}$ | | 5.13 $\times 10^{-60}$ | |

**Table S1: Organoid mass distributions.**
|  |  | 6 hours |  | 24 hours |  | 48 hours |  | 72 hours |  |
| --- | --- | --- | --- | --- | --- | --- | --- | --- | --- |
| BT-474 | | Mean mass ± SD | p-value | Mean mass ± SD | p-value | Mean mass ± SD | p-value | Mean mass ± SD | p-value |
| Vehicle | 1% DMSO | 1.12 ± 0.61 | - | 1.27 ± 0.69 | - | 1.52 ± 0.84 | - | 1.77 ± 1.00 | - |
| Staurosporine | 0.1 μM | 1.10 ± 0.57 | 1 | 1.22 ± 0.67 | 1 | 1.39 ± 0.82 | 0.0756 | 1.56 ± 0.96 | 0.0093 |
| | 1 μM | 1.02 ± 0.53 | 0.0901 | 0.76 ± 0.39 | 2.27 $\times 10^{-32}$ | 0.63 ± 0.34 | 1.28 $\times 10^{-57}$ | 0.57 ± 0.32 | 3.03 $\times 10^{-62}$ |
| | 10 μM | 1.02 ± 0.51 | 0.1342 | 0.79 ± 0.44 | 1.59 $\times 10^{-28}$ | 0.63 ± 0.36 | 4.55 $\times 10^{-56}$ | 0.56 ± 0.32 | 2.52 $\times 10^{-62}$ |
| Neratinib | 0.1 μM | 1.04 ± 0.46 | 1 | 0.97 ± 0.44 | 4.86 $\times 10^{-6}$ | 0.97 ± 0.45 | 5.09 $\times 10^{-14}$ | 0.98 ± 0.46 | 6.93 $\times 10^{-18}$ |
| | 1 μM | 1.08 ± 0.54 | 1 | 0.98 ± 0.49 | 4.34 $\times 10^{-7}$ | 0.86 ± 0.40 | 1.19 $\times 10^{-20}$ | 0.81 ± 0.36 | 1.48 $\times 10^{-27}$ |
| | 10 μM | 0.92 ± 0.42 | 0.0043 | 0.70 ± 0.31 | 3.73 $\times 10^{-21}$ | 0.59 ± 0.28 | 1.99 $\times 10^{-31}$ | 0.53 ± 0.21 | 4.71 $\times 10^{-34}$ |
| Lapatinib | 0.1 μM | 1.05 ± 0.52 | 1 | 1.13 ± 0.57 | 0.2512 | 1.23 ± 0.61 | 0.0025 | 1.31 ± 0.67 | 1.83 $\times 10^{-5}$ |
| | 1 μM | 1.08 ± 0.54 | 1 | 1.00 ± 0.49 | 3.37 $\times 10^{-5}$ | 0.96 ± 0.45 | 1.80 $\times 10^{-14}$ | 0.93 ± 0.43 | 1.70 $\times 10^{-21}$ |
| | 10 μM | 1.09 ± 0.47 | 1 | 0.91 ± 0.45 | 7.72 $\times 10^{-10}$ | 0.86 ± 0.40 | 3.31 $\times 10^{-20}$ | 0.83 ± 0.41 | 2.13 $\times 10^{-24}$ |
| Kruskal-Wallis p-value | | 0.0007 | | 4.25 $\times 10^{-70}$ | | 1.38 $\times 10^{-141}$ | | 7.68 $\times 10^{-168}$ | |

**Table S2.** Proportions of organoids that gained, lost, and maintained mass by treatment condition. (A) Proportions of MCF-7 organoids. (B) Proportions of BT-474 organoids. Data is plotted in Figure 5A.

|  |  |  |  | Organoid Behavior (%) |  |  |
| --- | --- | --- | --- | --- | --- | --- |
| MCF-7 |  | Time (h) | n | Gained Mass | Stable | Lost Mass |
| Vehicle | 1% DMSO | 12 | 800 | 23.2 | 67.4 | 9.4 |
|  |  | 24 | 794 | 44.8 | 36.0 | 19.1 |
|  |  | 48 | 764 | 48.6 | 24.1 | 27.4 |
|  |  | 72 | 715 | 50.8 | 16.6 | 32.6 |
| Staurosporine | 0.1 μM | 12 | 481 | 8.7 | 72.3 | 18.9 |
|  |  | 24 | 481 | 21.8 | 46.8 | 31.4 |
|  |  | 48 | 458 | 31.7 | 27.3 | 41.0 |
|  |  | 72 | 436 | 32.8 | 20.4 | 46.8 |
| | 1 μM | 12 | 508 | 10.2 | 74.6 | 15.2 |
|  |  | 24 | 505 | 11.7 | 56.6 | 31.7 |
|  |  | 48 | 490 | 13.7 | 36.9 | 49.4 |
|  |  | 72 | 460 | 10.0 | 31.3 | 58.7 |
| | 10 μM | 12 | 539 | 6.9 | 64.7 | 28.4 |
|  |  | 24 | 538 | 8.4 | 40.3 | 51.3 |
|  |  | 48 | 523 | 9.0 | 22.9 | 68.1 |
|  |  | 72 | 494 | 8.3 | 16.2 | 75.5 |
| Neratinib | 0.1 μM | 12 | 205 | 19.5 | 67.3 | 13.2 |
|  |  | 24 | 206 | 43.7 | 38.8 | 17.5 |
|  |  | 48 | 196 | 49.5 | 27.0 | 23.5 |
|  |  | 72 | 184 | 49.5 | 21.7 | 28.8 |
| | 1 μM | 12 | 222 | 18.0 | 71.6 | 10.4 |
|  |  | 24 | 221 | 37.6 | 45.2 | 17.2 |
|  |  | 48 | 214 | 44.4 | 30.8 | 24.8 |
|  |  | 72 | 203 | 46.3 | 24.6 | 29.1 |
| | 10 μM | 12 | 223 | 14.3 | 68.6 | 17.0 |
|  |  | 24 | 219 | 12.8 | 51.1 | 36.1 |
|  |  | 48 | 213 | 8.0 | 34.3 | 57.7 |
|  |  | 72 | 199 | 4.5 | 18.1 | 77.4 |
| Lapatinib | 0.1 μM | 12 | 218 | 21.1 | 66.5 | 12.4 |
|  |  | 24 | 214 | 35.5 | 43.5 | 21.0 |
|  |  | 48 | 203 | 42.9 | 24.6 | 32.5 |
|  |  | 72 | 187 | 43.3 | 22.5 | 34.2 |
| | 1 μM | 12 | 202 | 17.8 | 69.8 | 12.4 |
|  |  | 24 | 202 | 47.5 | 33.2 | 19.3 |
|  |  | 48 | 190 | 55.3 | 21.1 | 23.7 |
|  |  | 72 | 177 | 53.7 | 21.5 | 24.9 |
| | 10 μM | 12 | 251 | 15.5 | 62.5 | 21.9 |
|  |  | 24 | 252 | 32.5 | 31.0 | 36.5 |
|  |  | 48 | 238 | 36.1 | 18.9 | 45.0 |
|  |  | 72 | 220 | 37.7 | 10.0 | 52.3 |

|  |  |  |  | Organoid Behavior (%) |  |  |
| --- | --- | --- | --- | --- | --- | --- |
| BT-474 |  | Time (h) | n | Gained Mass | Stable | Lost Mass |
| Vehicle | 1% DMSO | 12 | 419 | 30.1 | 59.9 | 10.0 |
|  |  | 24 | 415 | 62.4 | 26.7 | 10.8 |
|  |  | 48 | 409 | 80.9 | 10.8 | 8.3 |
|  |  | 72 | 387 | 80.9 | 7.2 | 11.9 |
| Staurosporine | 0.1 μM | 12 | 335 | 31.0 | 55.2 | 13.7 |
|  |  | 24 | 334 | 49.7 | 38.3 | 12.0 |
|  |  | 48 | 334 | 65.9 | 17.7 | 16.5 |
|  |  | 72 | 327 | 72.8 | 11.0 | 16.2 |
|  | 1 μM | 12 | 277 | 11.9 | 39.0 | 49.1 |
|  |  | 24 | 276 | 10.9 | 19.9 | 69.2 |
|  |  | 48 | 275 | 5.5 | 12.0 | 82.5 |
|  |  | 72 | 266 | 2.6 | 9.4 | 88.0 |
|  | 10 μM | 12 | 276 | 11.6 | 47.5 | 40.9 |
|  |  | 24 | 273 | 12.1 | 24.9 | 63.0 |
|  |  | 48 | 273 | 6.6 | 13.9 | 79.5 |
|  |  | 72 | 266 | 4.5 | 7.9 | 87.6 |
| Neratinib | 0.1 μM | 12 | 121 | 14.9 | 57.0 | 28.1 |
|  |  | 24 | 120 | 15.0 | 40.8 | 44.2 |
|  |  | 48 | 121 | 20.7 | 34.7 | 44.6 |
|  |  | 72 | 116 | 28.4 | 22.4 | 49.1 |
|  | 1 μM | 12 | 132 | 12.9 | 56.8 | 30.3 |
|  |  | 24 | 131 | 15.3 | 37.4 | 47.3 |
|  |  | 48 | 131 | 10.7 | 22.1 | 67.2 |
|  |  | 72 | 128 | 7.0 | 22.7 | 70.3 |
|  | 10 μM | 12 | 103 | 12.6 | 42.7 | 44.7 |
|  |  | 24 | 106 | 8.5 | 30.2 | 61.3 |
|  |  | 48 | 105 | 4.8 | 8.6 | 86.7 |
|  |  | 72 | 103 | 0.0 | 7.8 | 92.2 |
| Lapatinib | 0.1 μM | 12 | 125 | 22.4 | 64.8 | 12.8 |
|  |  | 24 | 126 | 42.1 | 45.2 | 12.7 |
|  |  | 48 | 124 | 66.9 | 19.4 | 13.7 |
|  |  | 72 | 117 | 69.2 | 17.9 | 12.8 |
|  | 1 μM | 12 | 126 | 12.7 | 65.1 | 22.2 |
|  |  | 24 | 126 | 18.3 | 39.7 | 42.1 |
|  |  | 48 | 125 | 18.4 | 29.6 | 52.0 |
|  |  | 72 | 126 | 18.3 | 25.4 | 56.3 |
|  | 10 μM | 12 | 122 | 11.5 | 57.4 | 31.1 |
|  |  | 24 | 122 | 9.8 | 27.0 | 63.1 |
|  |  | 48 | 118 | 11.9 | 17.8 | 70.3 |
|  |  | 72 | 114 | 11.4 | 14.9 | 73.7 |

**Table S3:** Organoid viability analysis by endpoint ATP assay. Comparisons of cell viability measured by ATP assay. P-values calculated by unpaired t-test with Welch’s correction. Data is presented in Figure 5B.

|  |  | MCF-7 |  | BT-474 |  |
| --- | --- | --- | --- | --- | --- |
| | | Viability ± SD | p-value | Viability ± SD | p-value |
| Vehicle | 1% DMSO | 1.00 ± 0.22 | | 1.00 ± 0.22 | |
| Staurosporine | 0.1 μM | 0.50 ± 0.08 | <0.0001 | 0.78 ± 0.09 | 0.0006 |
| | 1 μM | 0.17 ± 0.04 | <0.0001 | 0.04 ± 0.01 | <0.0001 |
| | 10 μM | 0.13 ± 0.04 | <0.0001 | 0.01 ± 0.00 | <0.0001 |
| Neratinib | 0.1 μM | 1.02 ± 0.15 | 0.8463 | 0.40 ± 0.10 | <0.0001 |
| | 1 μM | 0.92 ± 0.12 | 0.4623 | 0.25 ± 0.04 | <0.0001 |
| | 10 μM | 0.08 ± 0.01 | <0.0001 | 0.13 ± 0.05 | <0.0001 |
| Lapatinib | 0.1 μM | 1.00 ± 0.04 | 0.9452 | 0.64 ± 0.11 | 0.0004 |
| | 1 μM | 1.01 ± 0.05 | 0.8598 | 0.38 ± 0.08 | <0.0001 |
| | 10 μM | 0.66 ± 0.08 | 0.0018 | 0.26 ± 0.03 | <0.0001 |
| | 50 μM | 0.14 ± 0.04 | <0.0001 | 0.01 ± 0.00 | <0.0001 |

### Supplementary Figures

**Figure S1:**
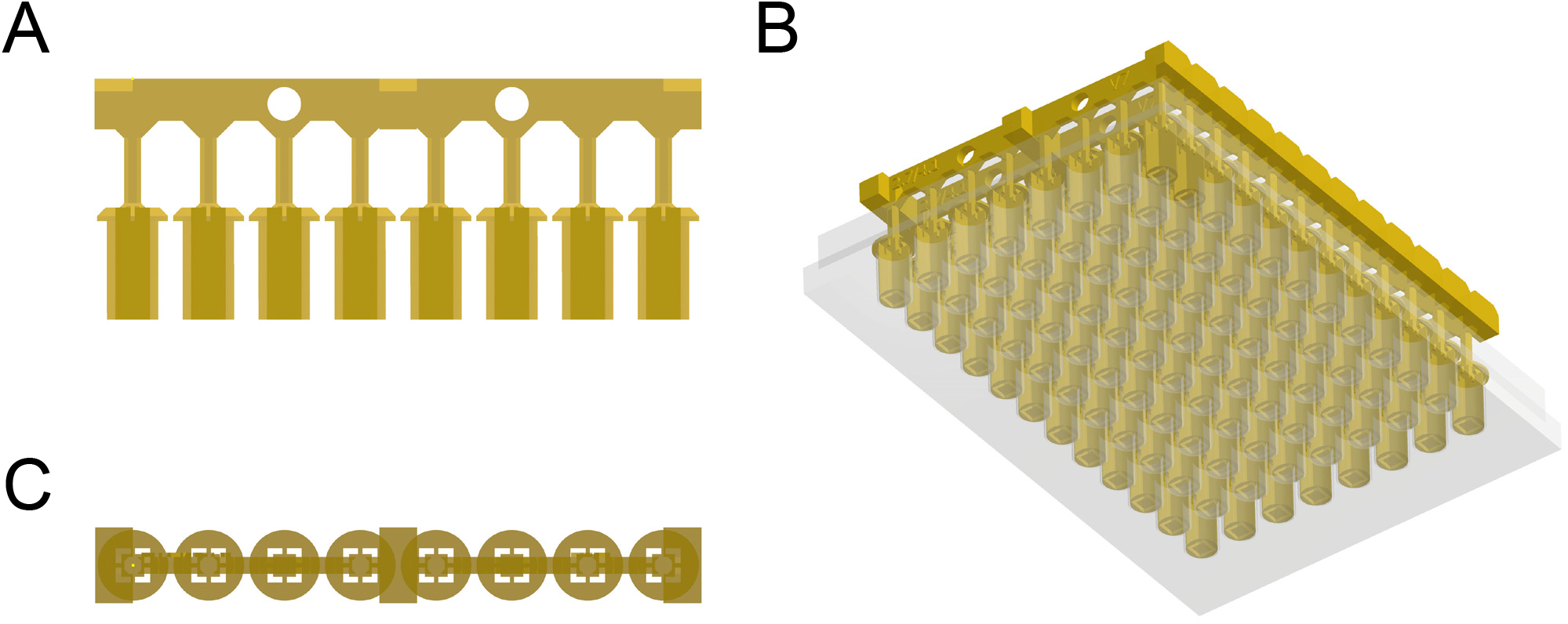
Schematics of well mask. (A) Side view. (B) Bottom view. (C) Plasma masks inserted into 96-well plate viewed from bottom.

**Figure S2:**
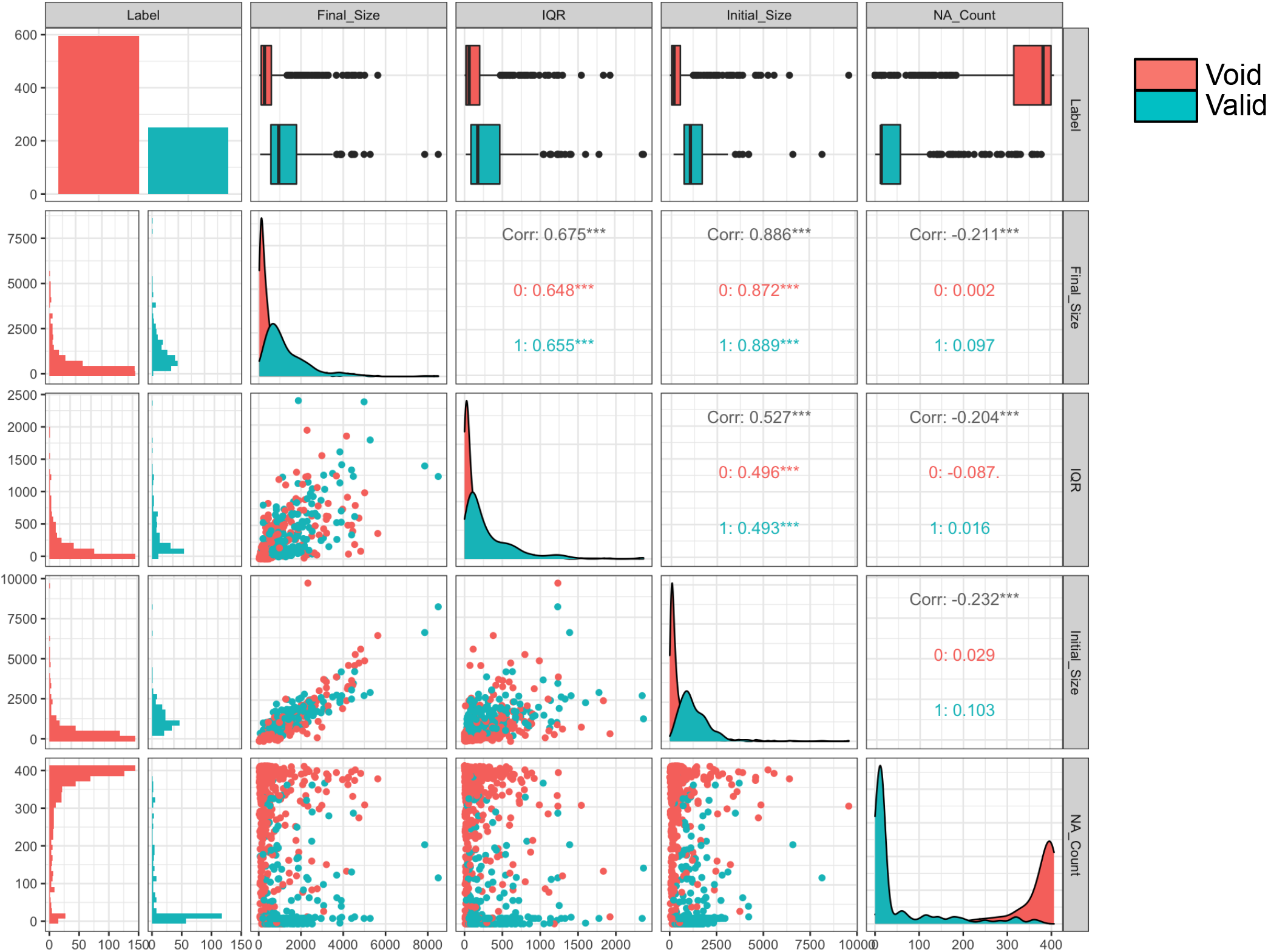
Pair-wise correlation matrix of classifier data. Extracted time-series analytical features of tumor growth patterns over recorded time. The number of missing frame (NA count), initial size, final size and interquartile range (IQR) was measured for each tracked tumor organoid. We labelled (n = 250 out of 846) tracked organoids as valid for downstream analysis. Pair-wise correlations are shown of the valid (label = 1) and void (label = 0) tracked organoids. Void tracks showed an increased number of missing frames and smaller IQR, initial and final size. Correlation of the classification data are shown within the paired subplots. Initial and final size were strongly correlated (R^2^ = 0.89). An XGBoost classifier was used to train a model to classify organoids as valid or void. We validated the model via cross-validation with 3-fold resampling of the sample population. The cross-validation score of the classifier was 91% with 93.5% accuracy.

**Figure S3:**
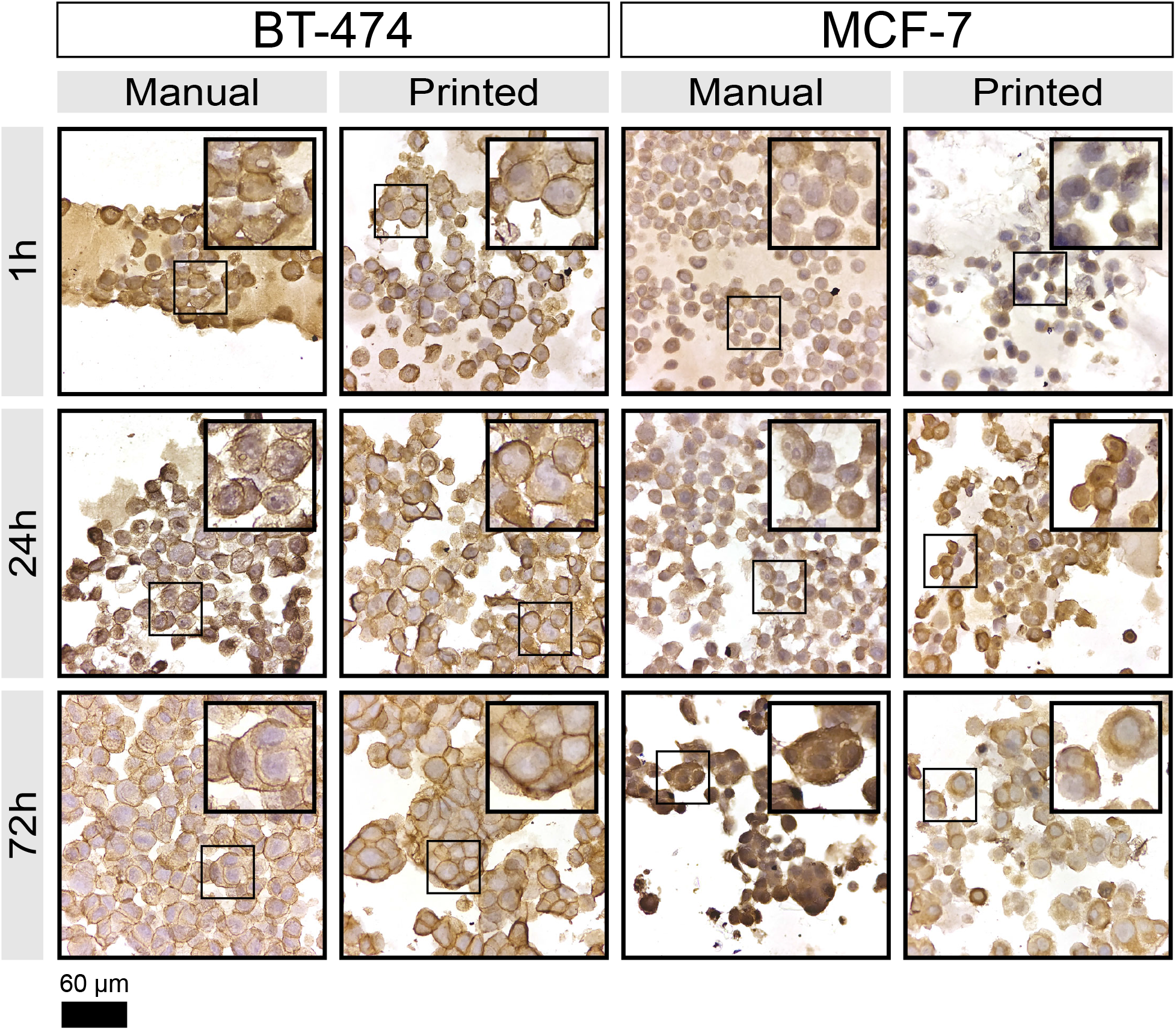
HER2 expression in BT-474 and MCF-7 organoids. Immunohistochemistry staining of 3D cultures for HER2. BT-474 cells have amplified HER2 expression^47,50^ while MCF-7 cells express lower levels of HER2 and lack HER2 amplification^47–49^.

**Figure S4:**
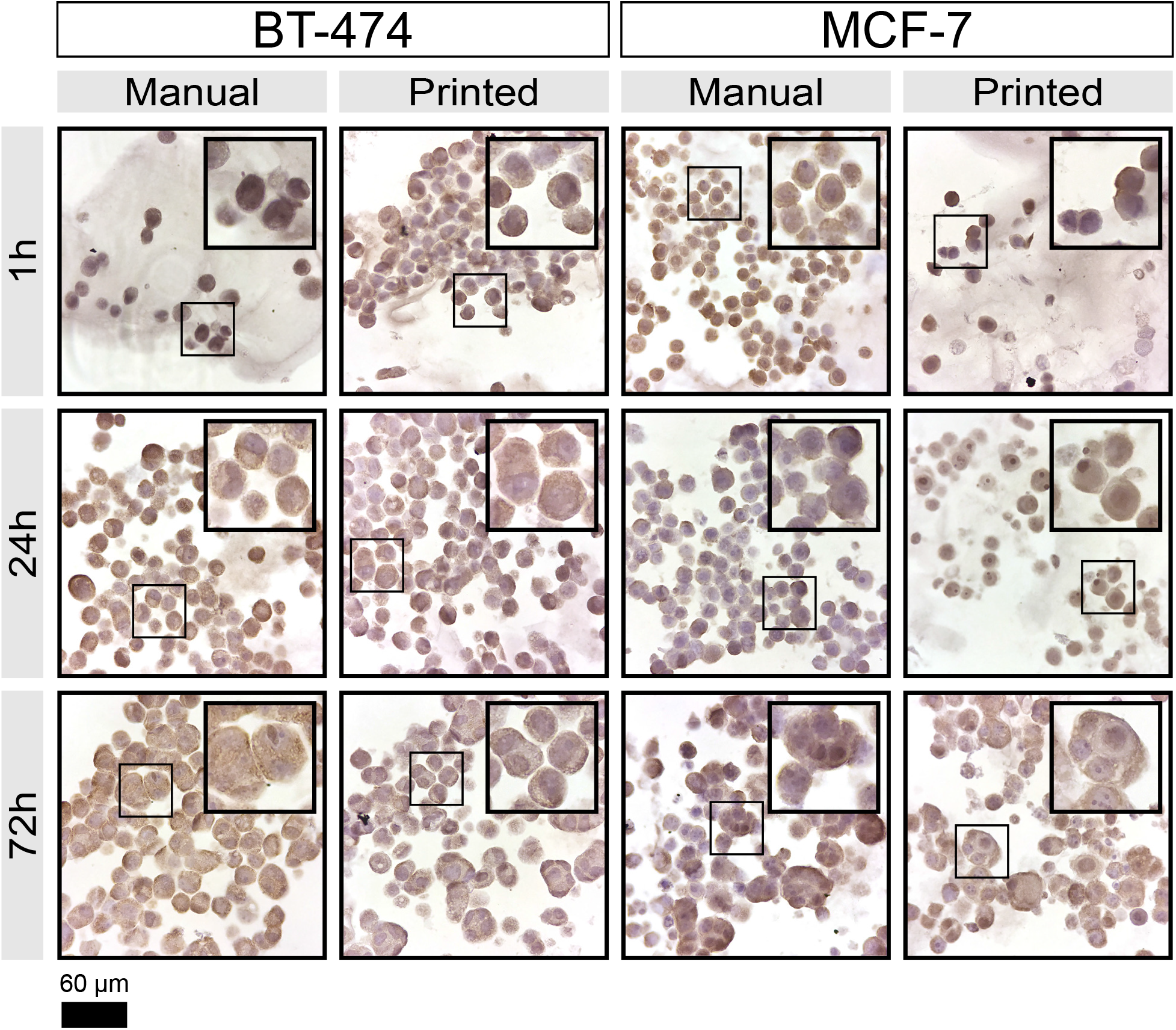
Estrogen Receptor Expression in BT-474 and MCF-7 organoids. Immunohistochemistry staining of 3D cultures for ER. Both BT-474 and MCF-7 cell lines are ER-positive^47–50^.

**Figure S5.**
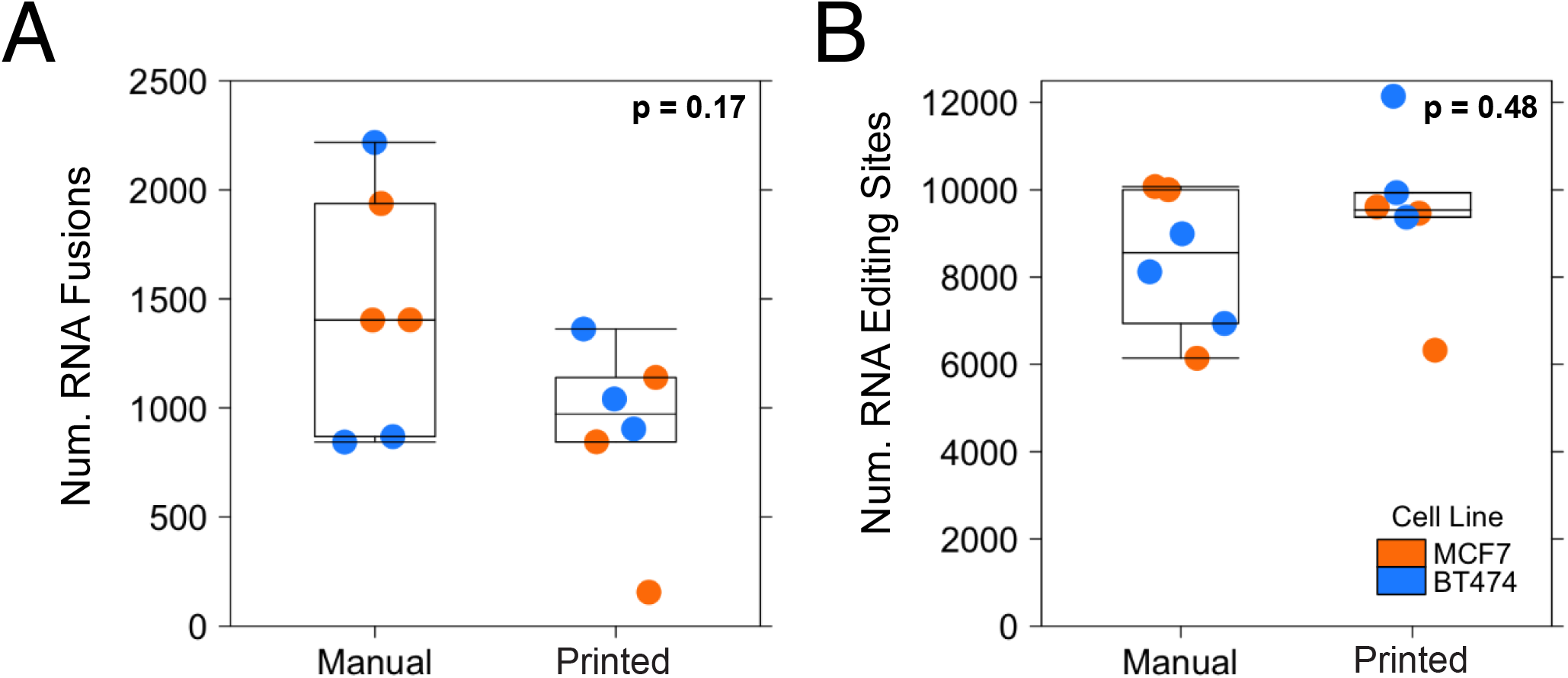
RNA Fusions and Editing Sites. (A) Number of RNA fusions detected by FusionCatcher by tumor organoid seeding method. The number of RNA fusions did not significantly differ between manually seeded and bioprinted organoids (p_BT-474_ = 0.179, p_MCF-7_ = 0.179). (B) The number of adenosine-to-inosine (A-to-I) RNA editing sites detected by REDItools were not associated with tumor organoid development method (p = 0.48, Mann-Whitney U-test). By cell line, the number of A-to-I RNA editing sites did not differ between printed and manually seeded organoids (p_BT-474_ = 0.1, p_MCF-7_ = 0.7).

**Figure S6:**
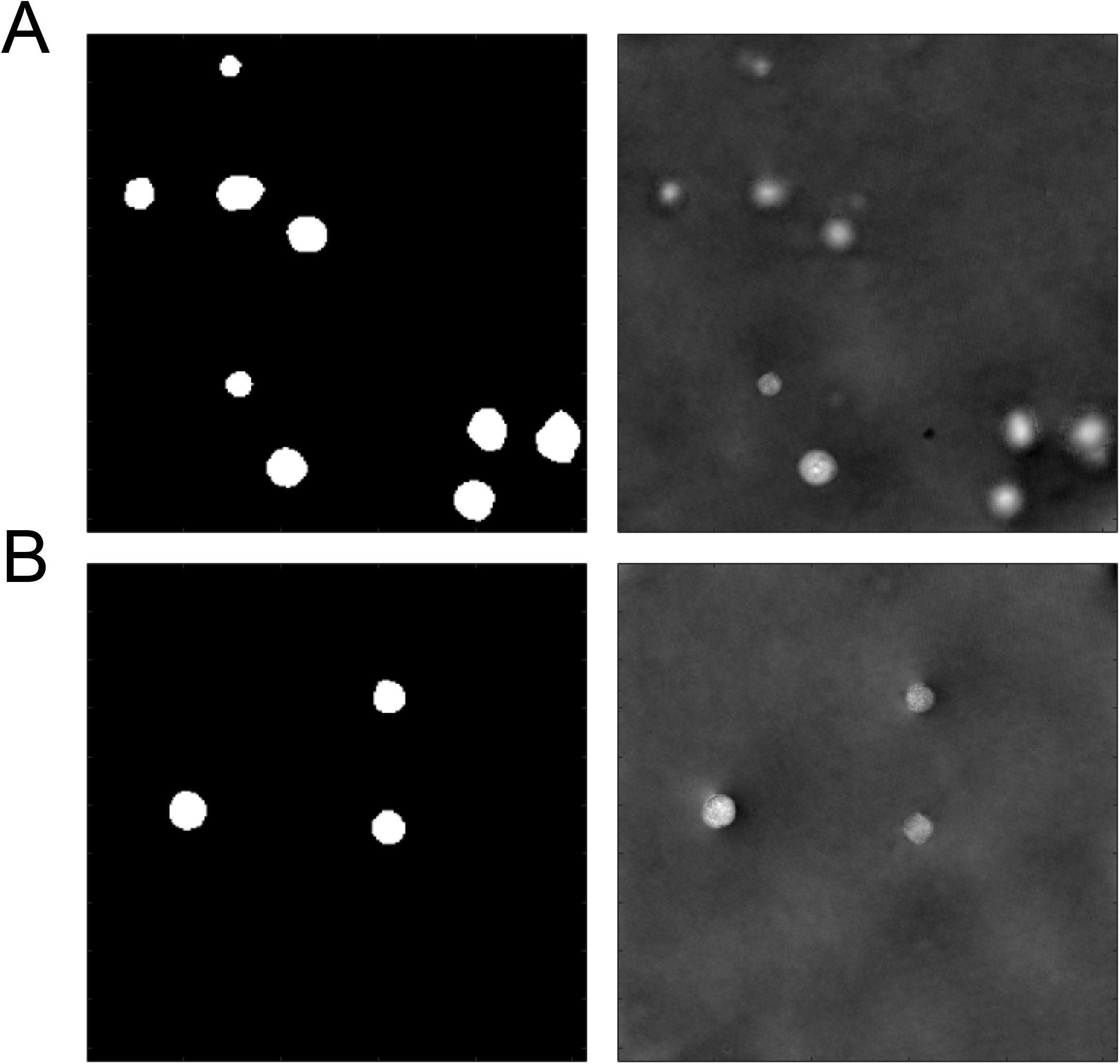
Image segmentation using a U-Net convolutional neural network. Representative masks (left) predicted by the U-Net-based segmentation algorithm for the background-corrected phase images (right).

**Figure S7:**
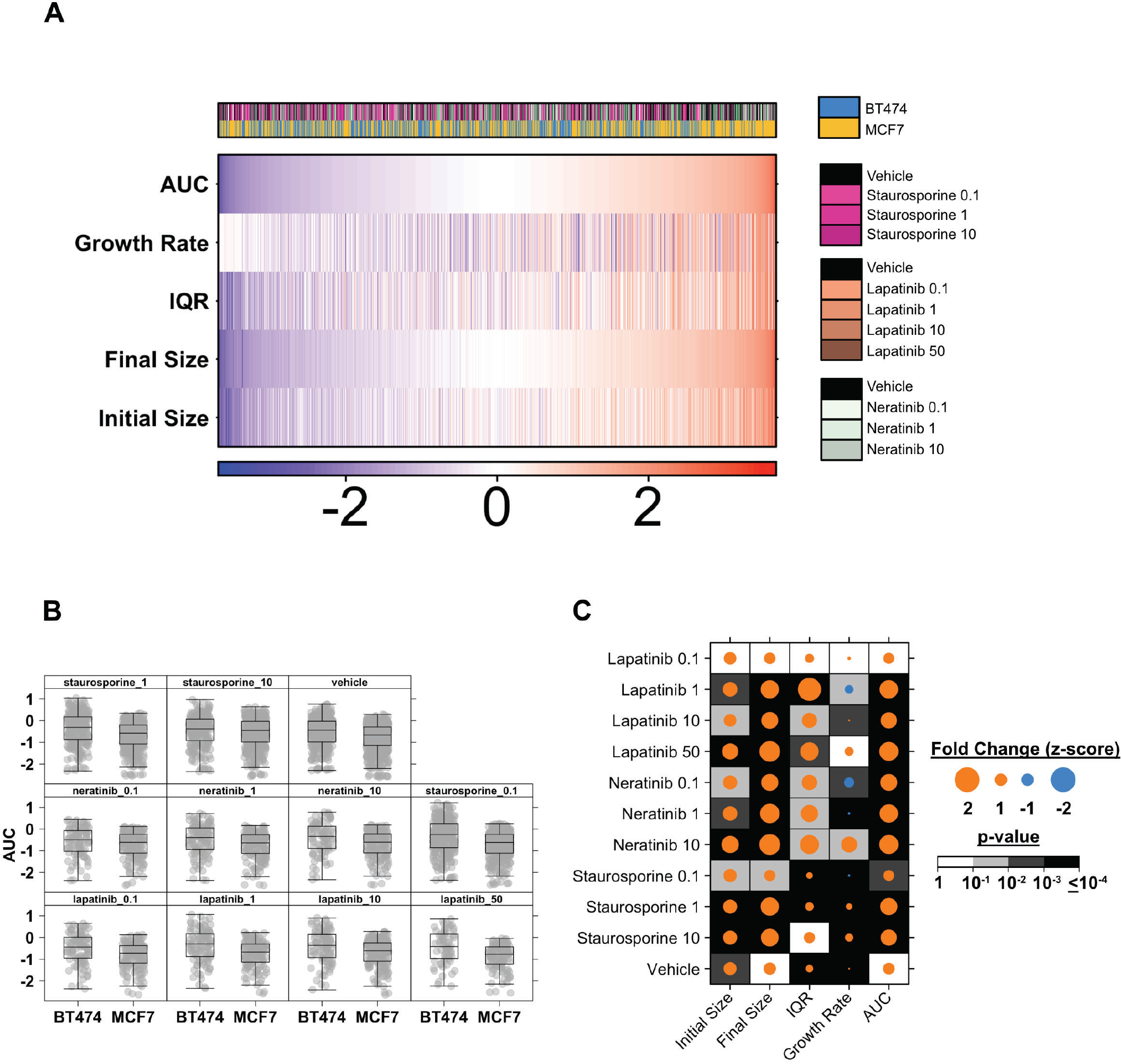
Growth patterns of MCF-7- and BT-474-derived tumor organoids among pharmacological treatments. We assessed the growth patterns of tumor organoids derived from MCF-7 and BT-474 breast cancer cell lines. Tumor organoids of both cell line types were grown in various pharmacological concentrations of lapatinib, neratinib and staurosporine ranging from 0.1 to 10 *μ*M. (A) Growth patterns of tumor organoids are arranged by area under the curve (AUC) metric measured by the integration of a fitted time-series step function. Z-transformed measurements of organoid AUC, linear growth pattern (R^2^ of a linear fit), initial size, final size and interquartile range (IQR) varied among sample population. (B) Overall growth patterns, measured as AUC, of MCF-7- and BT-474-derived organoids differed among each pharmacological treatment condition. (C) Fold change of growth patterns features were found to be significantly different among MCF-7 and BT-474 derived organoids in all three well treatment condition under various concentrations.

**Figure S8:**
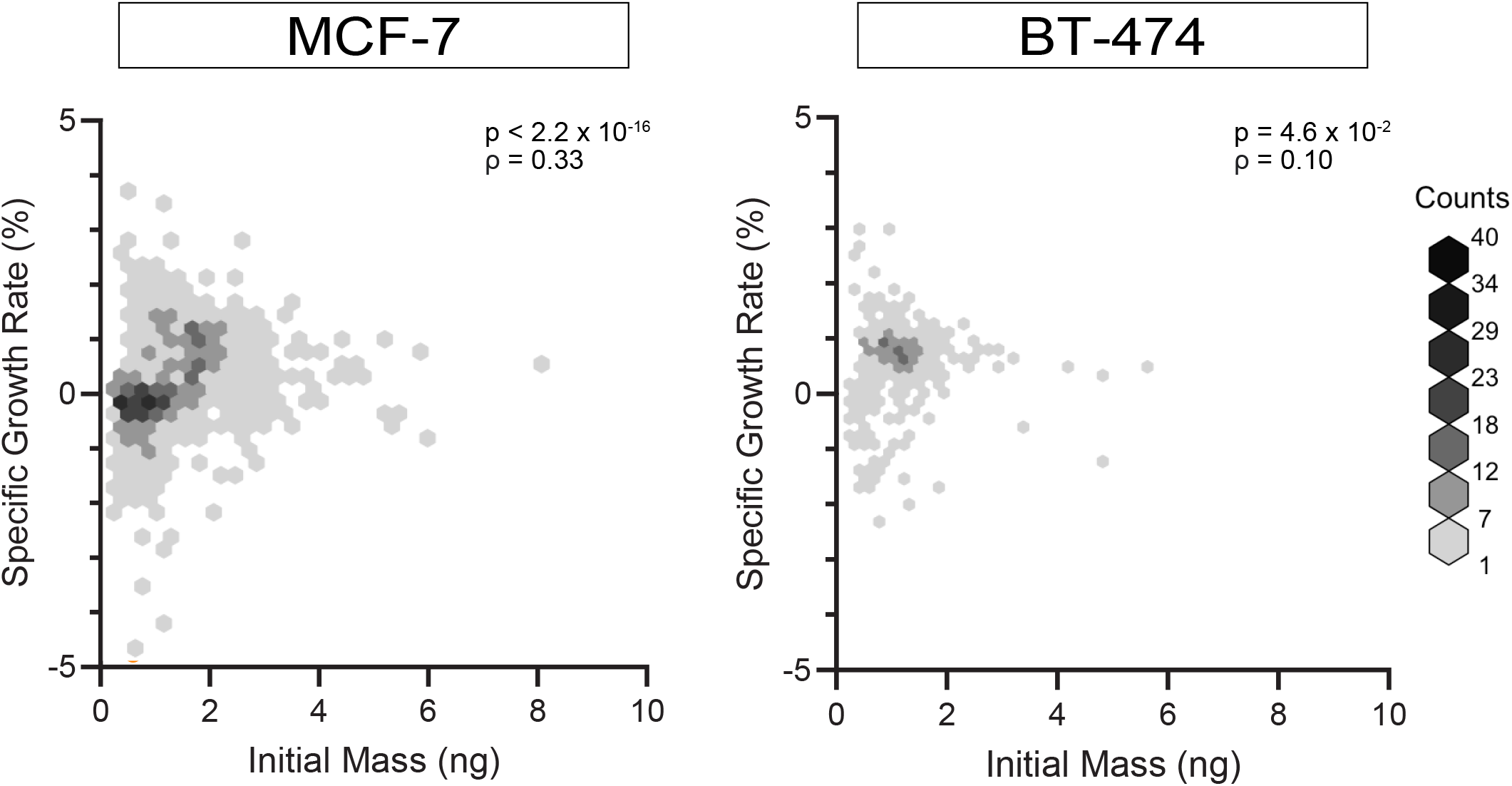
Specific growth rate correlates to initial organoid mass. Specific growth rate (growth in mass as a percentage of total mass) versus initial organoid mass was plotted for all organoids tracked. For both cell lines, we observe a positive relationship between initial organoid mass and specific growth rate. The association is stronger for MCF-7 organoids (Spearman’s ρ = 0.33, p < 2.2 x 10^-16^) compared to BT-474 organoids (ρ = 0.10, p = 4.6 x 10^-2^).

**Figure S9:**
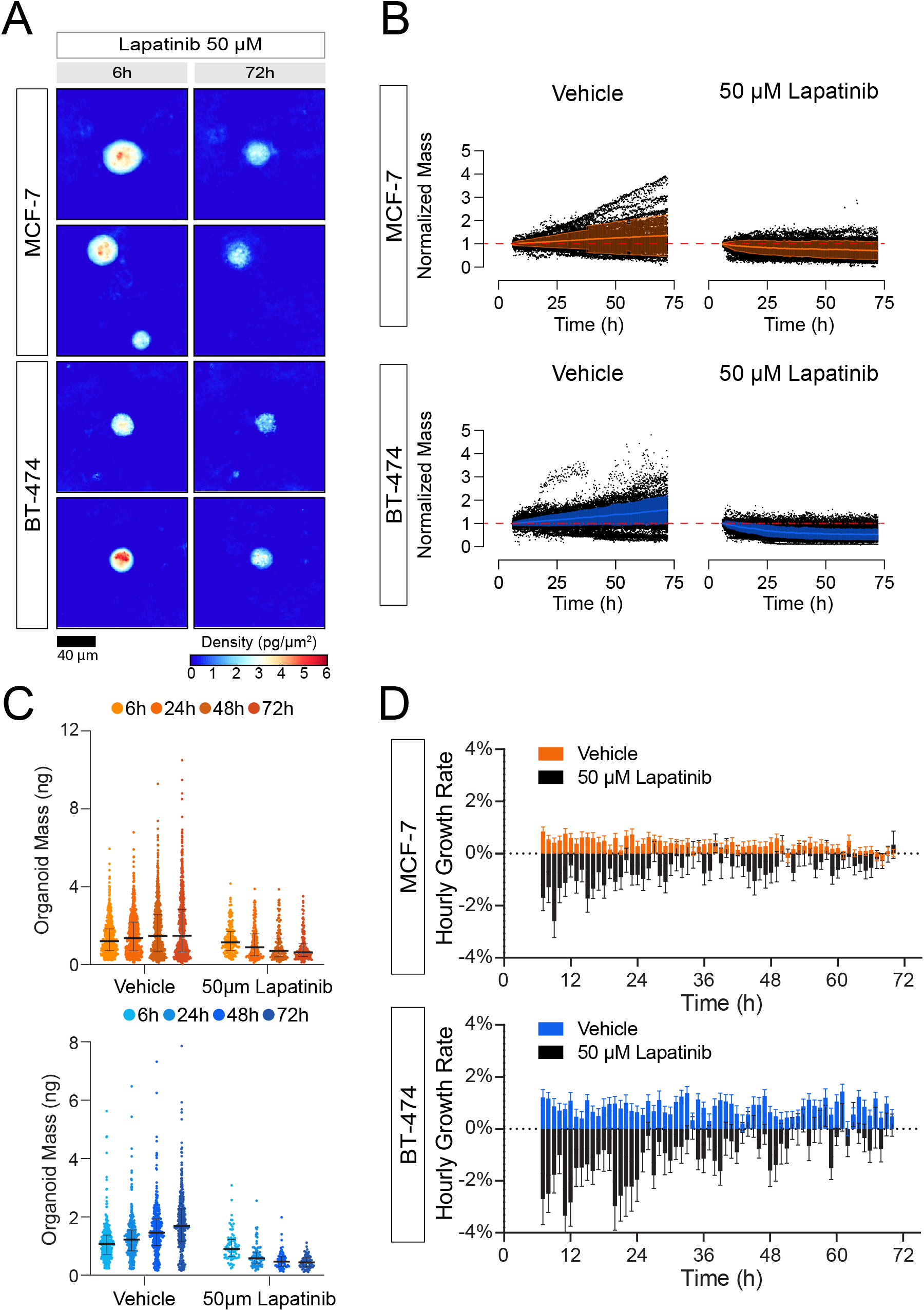
Response of bioprinted organoids to 50 *μ*M lapatinib. (A) Representative images of organoids treated with 50 *μ*M lapatinib. (B) 100 organoid tracks for are shown on each plot. The mean normalized mass ± standard deviation is also shown in orange (MCF-7) and blue (BT-474). (C) Mass distribution of tracked MCF-7 and BT-474 organoids by treatment. Each column represents the mass distribution at 6-, 24-, 28-, and 72-hours post-treatment (left to right). Black horizontal bars represent the median with error bars representing the interquartile range of the distribution. (D) Hourly growth rate comparisons (percent mass change) between organoids treated with 50 *μ*M lapatinib and vehicle.

**Figure S10:**
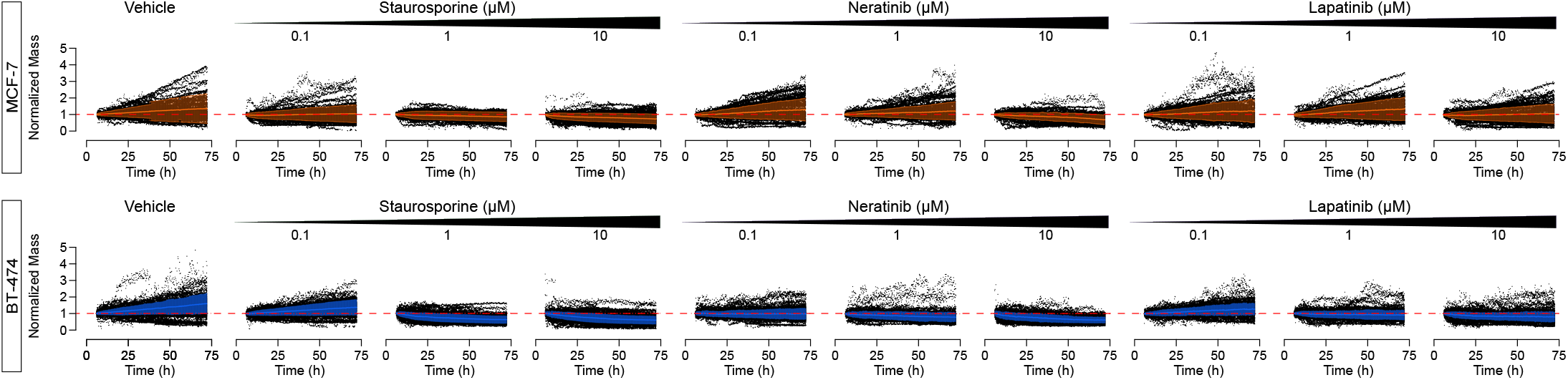
Representative normalized mass tracks by treatment condition. 100 organoid tracks for each treatment condition are shown on each plot. The mean normalized mass ± standard deviation is also shown in orange (MCF-7) and blue (BT-474).

**Figure S11:**
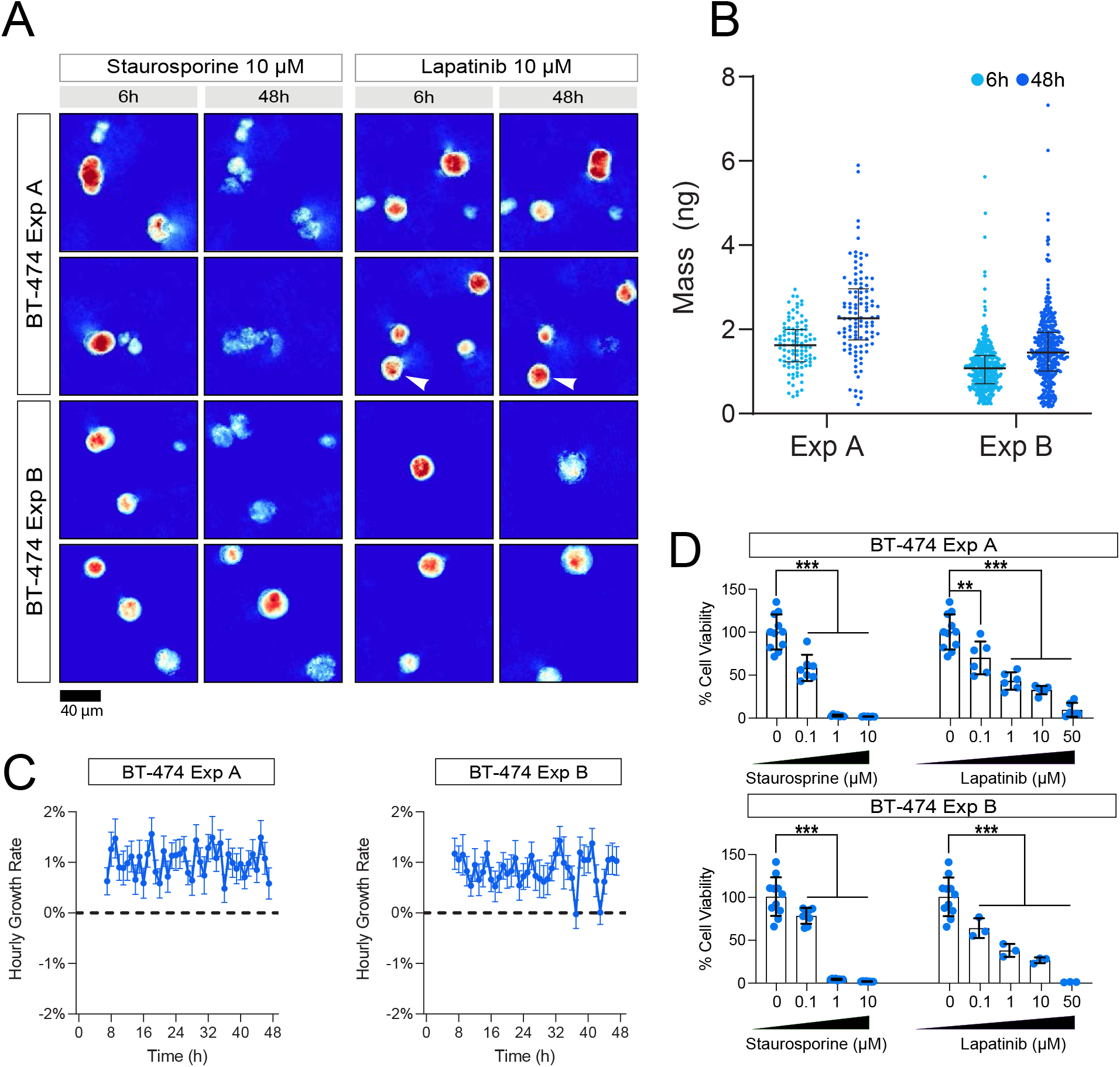
Comparison of BT-474 organoid datasets. Experiment A and B followed the same protocols with three exceptions. Experiment A was only imaged with the HSLCI for 48h hours after treatment, while Experiment B was imaged for 72 hours. Experiment A did not include neratinib in the drug screen. Experiment A was analyzed using our legacy pipeline as described in our original pre-print, while Experiment B was analyzed using the machine-learning based pipeline. (A) Representative images of organoids treated with 10 *μ*M staurosporine and 10 *μ*M lapatinib. (C) Mass distribution of tracked BT-474 organoids by treatment. The left column (pale blue) represents the mass distribution 6 hours post-treatment, while the right column (dark blue) represents the organoids 48 hours after treatment. Black horizontal bars represent the median with error bars representing the interquartile range of the distribution. (C) Hourly growth rate comparisons (percent mass change) between organoids treated with 50 *μ*M lapatinib and vehicle. (D) Percent cell viability of treated wells determined by an ATP-release assay. p<0.05 is denoted by *, p<0.01 is denoted by **, and p<0.001 is denoted by ***.

### Supplementary Results

**Mass reconstruction data extracted for classifier.** Mass reconstruction data of (n = 8,590) tumor organoid tracks derived from BT-474 and MCF-7 breast cancer cell lines were assessed for change in tumor size over 72 hours following treatment. Each cell line was treated with a series of concentrations of 0.1 to 50 *μ*M of vehicle (n = 1,593), lapatinib (n = 1,650), neratinib (n = 1,626) and staurosporine (n = 2,920). We assessed the organoid growth patterns using z-transformed measurements of area under the curve (AUC), linear growth rate, interquartile range, initial and final size (Supplementary Figure 8A). Sample population of tumor organoids from BT-474 and MCF-7 under various pharmacological treatments display a diverse set of growth patterns, arranged by AUC metric. Cell line and treatment type were dispersed across the sample population, thus supporting our decision to use a single set of classifier training data for both cell lines and all treatment conditions. We tested whether tumor organoids derived from BT-474 displayed differences in their growth patterns compared to organoids derived from MCF-7 (Supplementary Figure 8B). We found significant differences in growth patterns of BT-474 and MCF-7 derived organoids across treatments (Supplementary Figure 9C). In most conditions (10 out of 12 conditions), the initial size, final size and linear growth rate of MCF-7 derived organoids were larger than BT-474 organoids in all three pharmacological treatment conditions (Supplementary Figure 9B-C) (p < 0.0001, Mann-Whitney U-test). In two conditions, the linear growth rate of BT-474 derived organoids in 1 *μ*M lapatinib (fold change = −0.70, p = 2.76 x 10^-2^) and 0.1 *μ*M neratinib (fold change = −0.82, p = 3.82 x 10^-3^) were found to be greater than MCF-7 derived organoids. We did not find strong differences in growth patterns among BT-474 and MCF-7 derived tumor organoids in 0.1 *μ*M lapatinib treated wells.

